# Combinatorial Treatment of Glioblastoma with Temozolomide (TMZ) Plus 5-Ethynyl-2’-deoxyuridine (EdU)

**DOI:** 10.1101/2025.10.21.683081

**Authors:** Humeyra Kaanoglu, Yasemin K. Akyel, Adebimpe Adefolaju, Alain Valdivia, Dominique Higgins, Caroline Mosely, Corey J. Haswell, William C. Zamboni, Shawn D. Hingtgen, Andrew B. Satterlee, Aziz Sancar

**Affiliations:** Department of Biochemistry and Biophysics, University of North Carolina School of Medicine, Chapel Hill, North Carolina; Division of Pharmacoengineering and Molecular Pharmaceutics, Eshelman School of Pharmacy University of North Carolina, Chapel Hill, North Carolina; Department of Neurosurgery, University of North Carolina, Chapel Hill, North Carolina; UNC Advanced Translational Pharmacology and Analytical Chemistry Lab, UNC Eshelman School of Pharmacy, University of North Carolina at Chapel Hill, North Carolina; Eshelman Innovation, University of North Carolina, Chapel Hill, North Carolina

## Abstract

Glioblastoma (GBM) is the most aggressive malignant primary brain tumor in adults, with incidence peaking in later life. It is commonly treated with surgery followed by administration of ionizing radiation and the DNA alkylating agent temozolomide (TMZ). Even though this regimen confers some progression-free survival, there is essentially no cure with the median survival with the standard of care being about 12 months. Currently, several alternative approaches are being developed to improve upon this outcome. We have already shown EdU alone effectively treats GBM, and we now study efficacy of TMZ+EdU combination therapy. TMZ+EdU significantly improves antitumor efficacy compared to either single-agent therapy against GBM cell lines *in vitro*, against three different orthotopic GBM xenograft models, and against passage-zero patient GBM tumor tissues engrafted within an organotypic brain slice culture (OBSC)-based platform. Together, these data suggest that EdU could be effective alongside standard of care TMZ in patients with GBM.

**STATEMENT OF SIGNIFICANCE (50-WORD):** By simultaneously engaging distinct DNA repair pathways, combination of TMZ and EdU produced unprecedented survival benefits and synergistic effects in preclinical GBM models, offering a promising new avenue in the treatment of GBM.

## INTRODUCTION

Despite the application of multidisciplinary treatment approaches combining surgical, radiotherapeutic, and pharmacologic interventions, glioblastoma (GBM) remains one of the most treatment-resistant human cancers, with progression and recurrence as defining features (1–4). At the molecular level, GBM arises through a constellation of genetic alterations, including loss of heterozygosity on chromosome 10q, EGFR amplification, p16INK4a deletion, PTEN mutation, and, at later stages, TP53 and IDH1 mutations (5,6). Currently, the standard of care for GBM consists of surgery followed by ionizing radiation and temozolomide (TMZ) drug therapy (7–9)., During TMZ treatment, eventually MGMT (methylguanine methyl transferase) promoter hypermethylation causes silencing of MGMT which plays an essential role in TMZ-initiated death of malignant tumor cells. However, after an initial positive response to multimodal therapy, ∼80% of GBM patients experience recurrence at or near the original tumor site (10–13), often accompanied by new genetic alterations acquired under treatment pressure (14,15). At this stage, therapeutic options are limited, and even with standard of care, median survival remains around 12 months (4,13,16). Recently, a number of new drugs/intervention methods have been developed to improve the management of GBM (17). Among these, 5-ethynyl-2-deoxyuridine (EdU), conceptually, appears to be unique in terms of being immune to resistance development due to its high similarity to thymidine (see **Fig. 1** for the chemical structures of TMZ and EdU).

**Figure 1:**
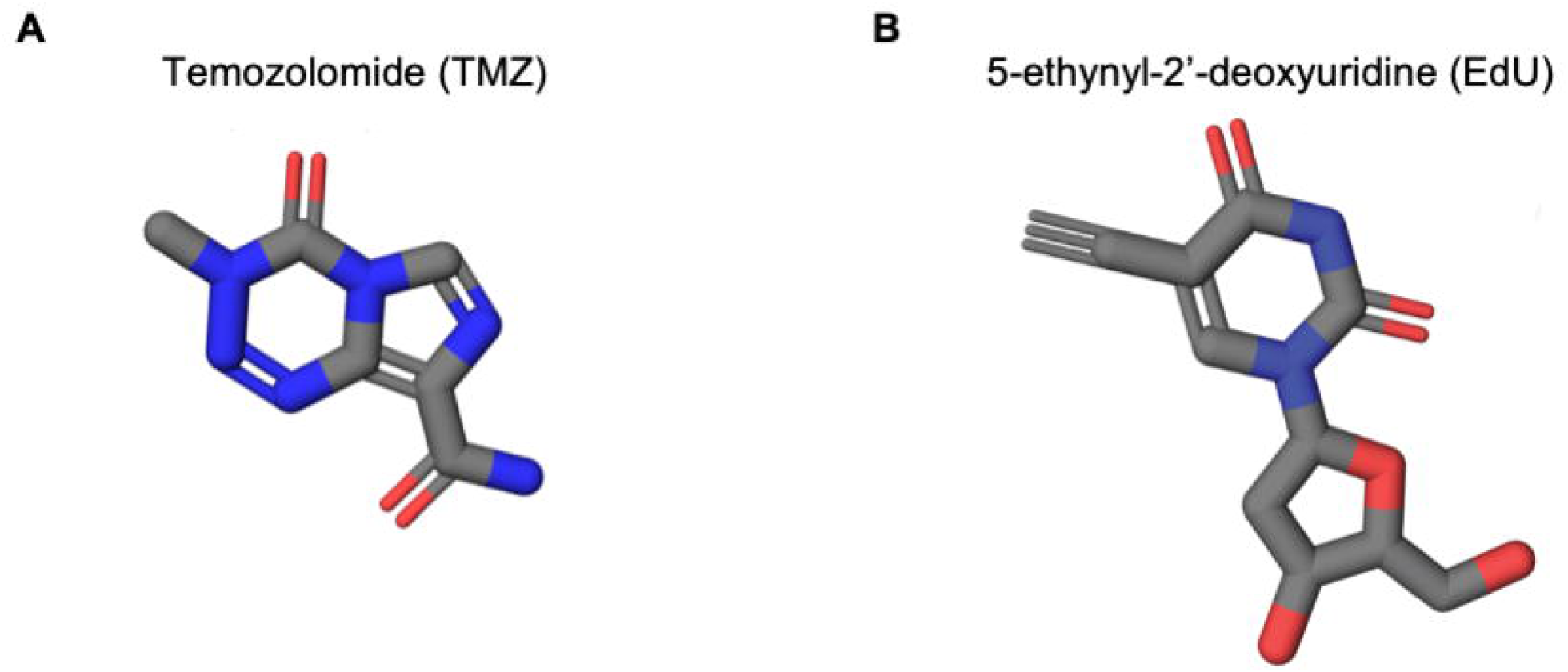
Chemical structures of temozolomide **(A)** and 5-ethynyl-2’-deoxyuridine (EdU) **(B)**.

EdU is commonly used to study DNA replication via "click chemistry" (18–21). During our attempt to employ EdU to analyze nucleotide excision repair patch size in human cell lines, we discovered that EdU incorporated into the human genome is recognized as “damage” and removed from the genome by the classic human dual incision mechanism in the form of 26-27 nt-long oligomers, leading to a futile cycle of incorporation into DNA and its removal (22). This realization, combined with previous reports on toxicity of EdU in tissue culture as well as its established property of crossing the blood brain barrier led to the notion that EdU might be effective against brain cancer (23,24). In our previous study, we showed that EdU is effective against GBM cell lines *in vitro*, orthotopic *in vivo* mouse models of GBM established with intracranial engrafts of mouse glioma and human GBM cells, and living, passage-zero patient GBM tissues tested *ex vivo* on an organotypic brain slice culture (OBSC)-based platform (25). Here, building on these proof-of-concept studies, we sought to investigate the therapeutic potential of combining TMZ with EdU for GBM, reasoning that their distinct mechanisms of action and the conceptual improbability of resistance to EdU—given its close structural similarity to thymidine—might reduce recurrence rates and even enable durable remission in a subset of patients. To this end, in this study, we tested the synergistic effect of TMZ+EdU in three different GBM cell lines *in vitro*, in *in vivo* orthotopic mouse models of GBM established with these three GBM cell lines, and in living, passage-zero GBM patient tissues seeded on OBSCs. Our results show the combination of TMZ and EdU as a new and promising therapeutic avenue, yielding synergistic effects in GBM and having the potential to ultimately improve the poor prognosis faced by patients with this disease.

## RESULTS

### Targeted localization and pharmacokinetic features of EdU *in vivo*

In our previous study, we showed the targeted localization of EdU into the human GBM cell lines seeded on OBSCs (25). In this study, we started out by testing the targeted localization of EdU into tumor cells *in vivo*. To this end, we established orthotopic GBM tumors in female athymic nude mice by inoculating FLuc expressing U87 human GBM cells into their brain parenchyma. Since U87 tumor cells form large and well-demarcated tumors upon intracranial inoculation, we chose this *in vivo* orthotopic mouse model of GBM to test the targeted localization of EdU into tumor cells in the brain (26,27). Upon confirming tumor establishment via bioluminescent imaging (BLI), we randomized these mice into four different groups to be injected with PBS Control, 200 mg/kg EdU, 5 mg/kg TMZ, or 5 mg/kg TMZ + 200 mg/kg EdU. These injections were administered for 3 consecutive days, and the brains of mice were harvested the following day to perform hematoxylin and eosin (H&E), nuclear (DAPI) and EdU staining. In **Figure 2**, the presence of tumor mass is seen with H&E staining. Once the tumor cells were confirmed with H&E, EdU staining via click chemistry along with nuclear staining (DAPI) was performed on the slices from the same area of the brains. As negative controls, there was no EdU signal detected in either tumor or non-tumor cells in the brains of mice administered with PBS or 5 mg/kg TMZ, pointing out the specificity of EdU staining via click chemistry (**Fig 2A, 2B, 2E and 2F**). On the other hand, the EdU signal (green) was highly localized to the tumor cells in mice treated with both 200 mg/kg EdU (**Fig. 2C**) and 5 mg/kg TMZ + 200 mg/kg EdU (**Fig. 2D**). The quantification of EdU-positive cells in tumor versus non-tumor cells revealed ∼83 times more EdU-positive cells in 200 mg/kg EdU-treated group and ∼70 times more EdU positive cells in 5 mg/kg TMZ and 200 mg/kg EdU-treated group (**Fig. 2G, ****p<0.0001 for both comparisons**). The residual EdU signal observed in **Fig. 2C-2D** was localized mostly to the subventricular zone (SVZ) of the brain, which is known to harbor some proliferative cells (28). These results confirmed the highly targeted localization of EdU into tumor cells *in vivo*. To our knowledge, EdU’s distribution in blood plasma and brain has not been characterized before. To characterize this biodistribution and clearance of EdU *in vivo*, we next performed a pharmacokinetic study with healthy female athymic nude mice. We administered a single dose of 200 mg/kg EdU, collected blood plasma and brain samples from mice at six different time points ranging from 15 mins to 6 hrs, and obtained the EdU concentration over time via LC-MS/MS (**Fig. 3A**). In plasma, the maximum concentration of EdU was reached at 15 minutes at ∼73 μg/ml, with a half-life of ∼10 minutes (**Fig. 3B**). On the other hand, in brain, the maximum concentration of EdU was reached at 15 minutes at ∼4.8 μg/ml, with a half-life of ∼30 minutes (**Fig. 3C**). The mean AUC_0–∞_ values for plasma and brain were 42.14 and 4.92, respectively, corresponding to a brain-to-plasma (B/P) AUC ratio of approximately 0.12 for EdU. Together, these results supported our hypothesis that EdU gets specifically localized into the tumor cells in the brain *in vivo* and identified the pharmacokinetic parameters of EdU in blood plasma and brain tissue for the first time.

**Figure 2:**
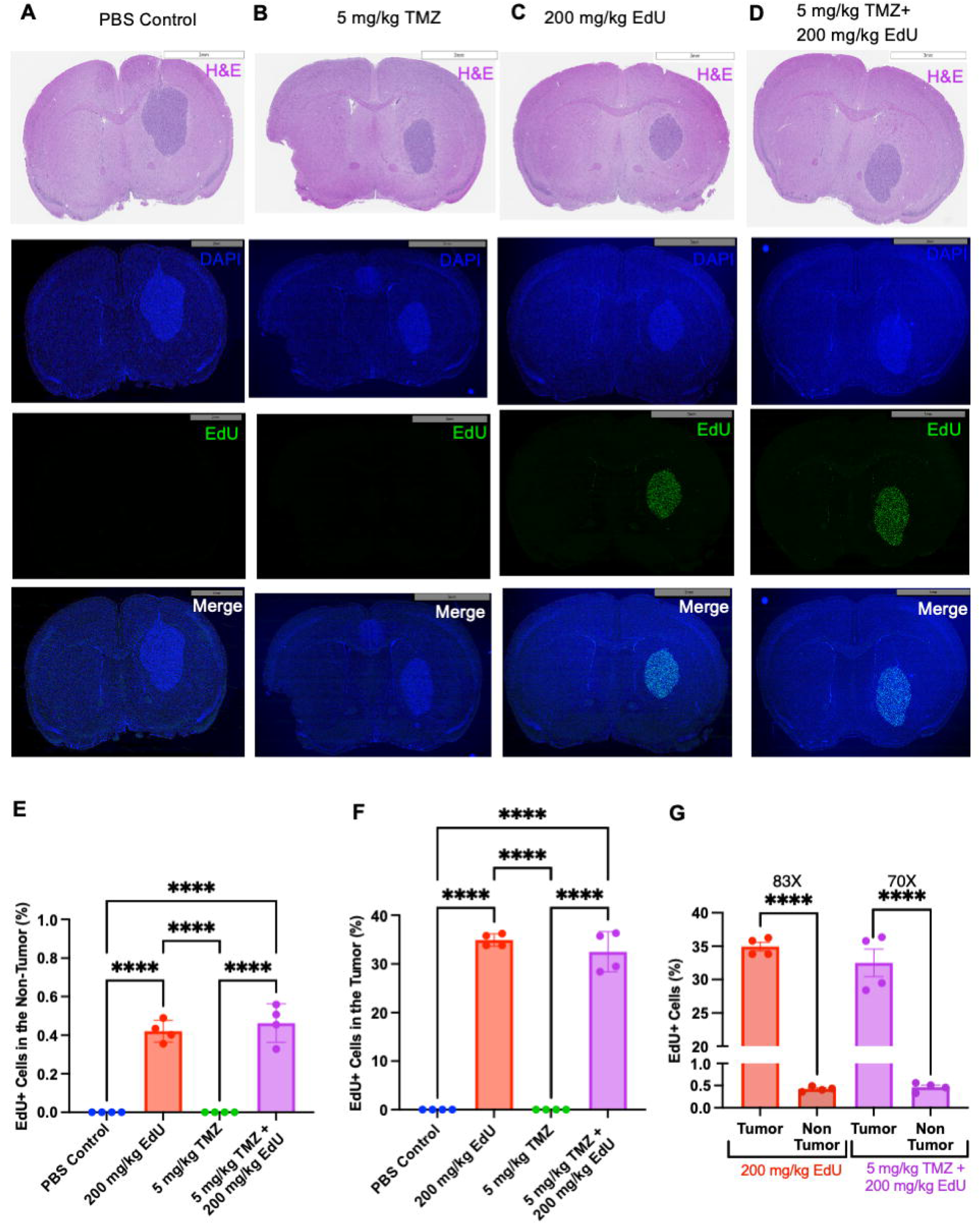
Targeted localization of EdU into GBM tumor cells *in vivo* in single-agent EdU- and TMZ+EdU combination-treated mice. Hematoxylin and eosin (H&E), nuclear (DAPI), EdU staining, and merged DAPI and EdU staining pictures in the brains of PBS-**(A)**, 5 mg/kg TMZ-**(B)**, 200 mg/kg EdU-**(C)**, and 5 mg/kg TMZ + 200 mg/kg EdU-treated **(D)** mice bearing orthotopic xenografts of U87. **(E)** Quantification of EdU-positive cells in non-tumor cells in the brains of PBS-(n=4), 200 mg/kg EdU-(n=4), 5 mg/kg TMZ-(n=4), and 5 mg/kg TMZ + 200 mg/kg EdU-treated (n=4) mice bearing orthotopic xenografts of U87 (One-way ANOVA, ****p<0.0001). **(F)** Quantification of EdU-positive cells in tumor cells in the brains of PBS-(n=4), 200 mg/kg EdU-(n=4), 5 mg/kg TMZ-(n=4), and 5 mg/kg TMZ + 200 mg/kg EdU-treated (n=4) mice bearing orthotopic xenografts of U87 (One-way ANOVA, ****p<0.0001). **(G)** Quantification of EdU-positive cells in tumor versus non-tumor cells in the brains of 200 mg/kg EdU-(n=4) and 5 mg/kg TMZ + 200 mg/kg EdU-treated (n=4) mice bearing orthotopic xenografts of U87 (Unpaired, two-tailed t-test, ****p<0.0001).

**Figure 3:**
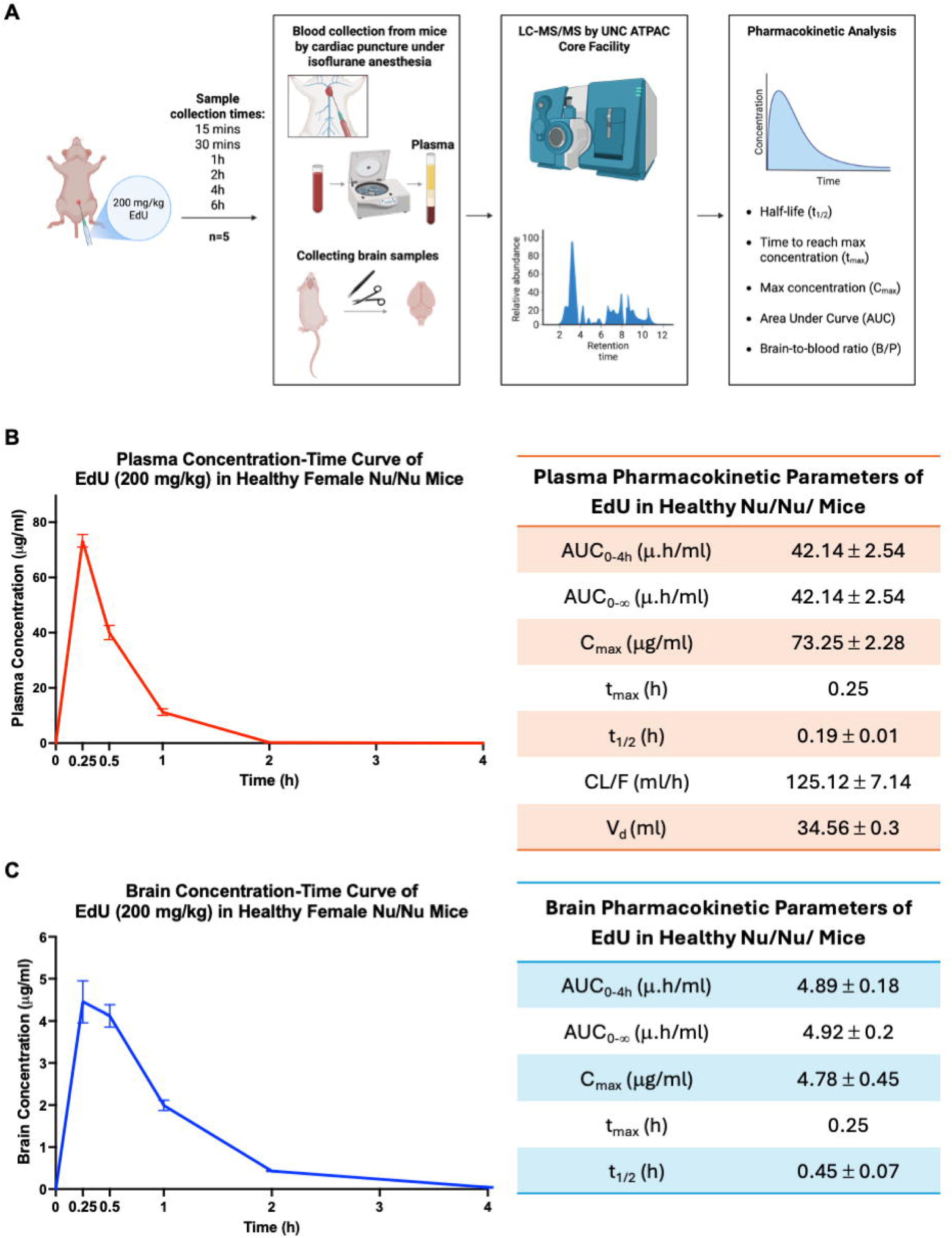
Pharmacokinetic features of EdU in blood plasma and brain. **(A)** Overview of the sample collection and analysis to identify pharmacokinetic parameters of EdU in blood plasma and brain of female athymic nude mice. **(B)** EdU concentration versus time curve in blood plasma, and Area Under Curve (AUC), maximum concentration (C_max_), time to reach maximum concentration (t_max_), half-life (t_1/2_), apparent clearance (CL/F) and volume of distribution (V_d_) parameters for EdU in blood plasma. **(C)** EdU concentration versus time curve in brain, and Area Under Curve (AUC), maximum concentration (C_max_), time to reach maximum concentration (t_max_), and half-life (t_1/2_) parameters for EdU in brain.

### Testing TMZ+EdU combination therapy *in vitro* in human GBM cells and *in vivo* in orthotopic mouse models of GBM

It has been found that combinations of various mutations in oncogenes and tumor suppressor genes give rise to GBM (5). Human GBM cell lines derived from patients are widely used in testing drug candidates both *in vitro* and *in vivo*. To recapitulate this heterogeneous genetic profile, in our study, we used 3 human GBM cell lines with different sets of mutations (27,29–34) in testing the combination of TMZ and EdU: U87, GBM8, and LN229. All of these cell lines are engineered to express firefly luciferase (FLuc) and thus bioluminescence signal measured upon incubation with luciferin is used as a measure of viability. We tested these cell lines with the standard of care drug TMZ and EdU, and the TMZ+EdU combination. Testing was done by cell survival assay *in vitro* and by monitoring survival and tumor growth *in vivo* in female athymic nude mice bearing orthotopic xenografts of these 3 cell lines.

### Testing TMZ+EdU combination therapy in vitro in U87 cell line and in mice bearing orthotopic xenografts of U87

We first tested for a potential synergistic effect of TMZ and EdU combination on U87 cells *in vitro* before proceeding to *in vivo* with this cell line. We have reported the IC50 value of EdU for this cell line as 40 μM (25). Cell survival assay with the combination of EdU at this concentration and various doses of TMZ showed that these two drugs are synergistic, especially at the higher doses of TMZ (**Fig. S1A**). This *in vitro* synergism between TMZ and EdU encouraged us to test this combination *in vivo* in the mice bearing orthotopic xenografts of U87 cells.

To establish the *in vivo* orthotopic mouse model with the U87 cell line, we inoculated U87 cells intracranially into the brain parenchyma of female athymic nude mice and confirmed establishment of tumors via BLI. Informed by our results on the pharmacokinetic features of EdU, we decided to administer EdU at 200 mg/kg every Monday-Friday at a once daily frequency and TMZ at 5 mg/kg every Monday, Wednesday, and Friday once every other day for a total of 6 weeks. For the combination group, mice received the TMZ injections in the morning and EdU injections in the afternoon with the indicated regimens for 6 weeks. Upon randomization of mice into PBS control and three treatment groups on Day 7 (**Fig. S1B**), we monitored the tumor growth of these mice via BLI. Neither 200 mg/kg EdU nor 5 mg/kg TMZ treatment showed a successful suppression of tumor growth while the treatment with the combination of 200 mg/kg EdU and 5 mg/kg TMZ suppressed the tumor growth and decreased the tumor signal below the baseline level (**Fig. 4A**). This suppression lasted as long as the injections continued. However, once the injections were completed by Day 46, the tumors recurred in 7-10 days, and the mice started to reach humane end point after ∼15 days following the discontinuation of the treatment. **Figure 4B** shows the BLI images of representative mice throughout the study (see **Fig. S1B-G** for extended BLI images). While treatment with single-agent 200 mg/kg and 5 mg/kg TMZ extended the survival by ∼30% and ∼50%, respectively, compared to the PBS control group (p<0.0008 and p<0.0001, respectively), combination of 5 mg/kg TMZ and 200 mg/kg EdU extended the survival by more than 130% compared to PBS control group (p<0.0001). Additionally, combination group extended the survival by ∼60% and ∼80% compared to the single-agent TMZ and EdU groups, respectively (**Fig. 3C**, p<0.0001 for both comparisons). At the study endpoint of this study, 1 out of 8 mice in the combination group showed no detectable tumor BLI signal, and the brain of this mouse did not show detectable tumor cells at or around the tumor cell inoculation site (**Fig. S1G-H**). Moreover, at the study endpoint, ∼%12 of mice (1 out of 8 mice) in 5 mg/kg TMZ + 200 mg/kg EdU group was alive. Thus, we concluded that TMZ and EdU combination showed a synergistic effect *in vivo* in suppressing the tumor growth and increasing the survival in mice bearing orthotopic xenografts of U87 cells.

**Figure 4:**
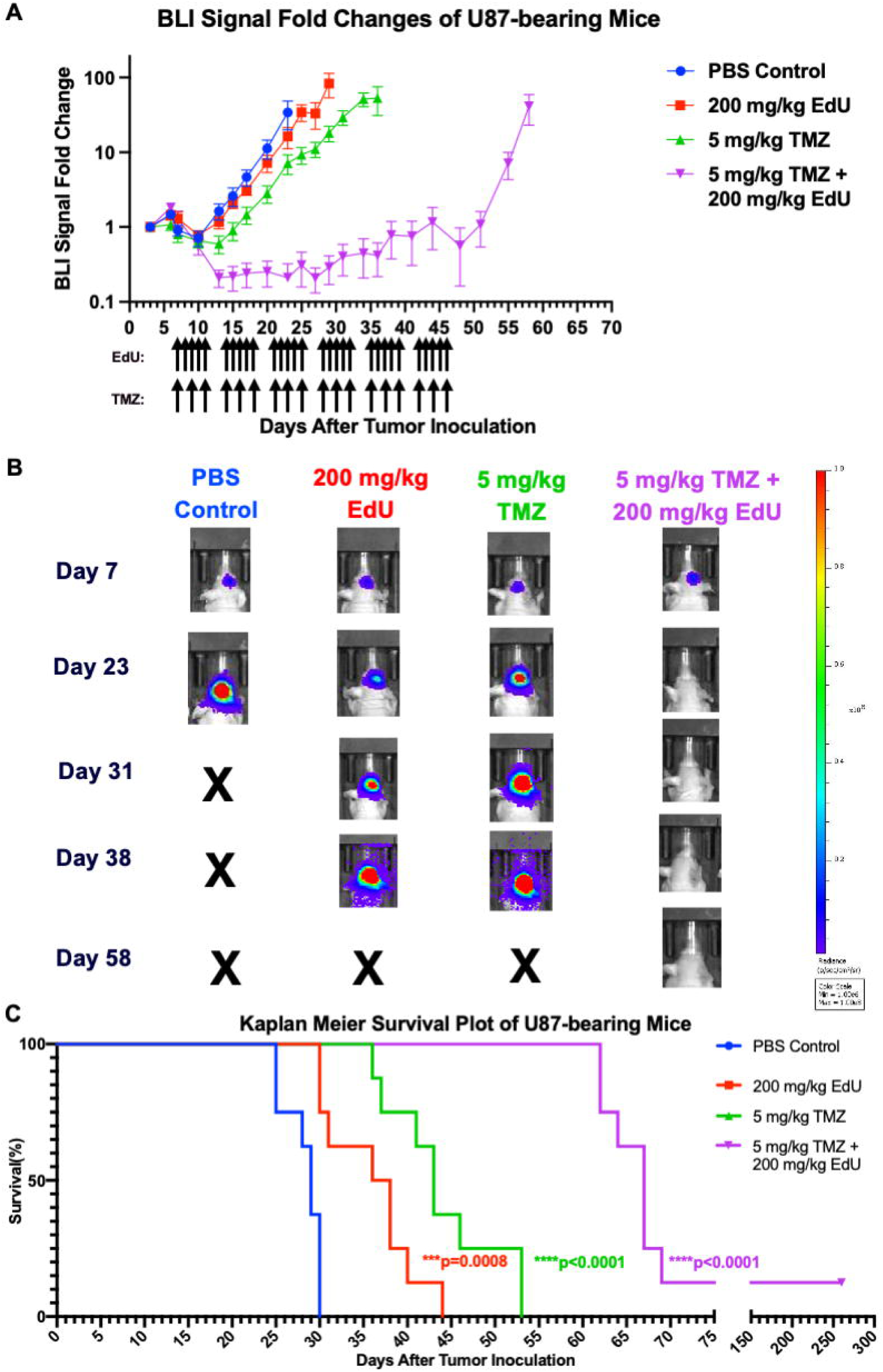
Testing the efficacy of TMZ+EdU in *in vivo* orthotopic U87 mouse model of GBM. **(A)** Bioluminescence imaging (BLI) signal fold changes of the tumors of mice bearing orthotopic xenografts of U87 and treated with either PBS (n=8), 200 mg/kg EdU (n=8), 5 mg/kg TMZ (n=8), or 5 mg/kg TMZ + 200 mg/kg EdU (n=8). Arrows under the x axis denote the days on which six weeks of EdU and TMZ injections were administered. **(B)** Representative BLI images of mice treated with either PBS, 200 mg/kg EdU, 5 mg/kg TMZ, or 5 mg/kg TMZ + 200 mg/kg EdU on Days 7, 23, 31, 38, and 58. **(C)** Kaplan Meier survival plot of mice bearing orthotopic xenografts of U87 and treated with either PBS, 200 mg/kg EdU, 5 mg/kg TMZ, or 5 mg/kg TMZ + 200 mg/kg EdU (log-rank test).

### Testing the synergistic effect of TMZ and EdU combination in vitro in the GBM8 cell line and in mice bearing orthotopic xenografts of GBM8

Upon obtaining these results with the U87 *in vivo* model, we moved on to test the potential synergistic effect of the combination of TMZ and EdU on GBM8 cell line *in vitro*. Previously, the IC50 value of EdU for this cell line was identified as ∼50 nM (25). Cell survival assay combining this IC50 value of EdU with various doses of TMZ showed synergistic to additive effect at the high doses of TMZ (**Fig. S2A**). Although the synergism observed in GBM8 was not as high as that in U87 in cell survival assay, it was encouraging enough to proceed with GBM8 cell line to establish an *in vivo* orthotopic mouse model of GBM.

To this end, we started out by inoculating GBM8 cells intracranially into the brain parenchyma of female athymic nude mice. After confirmation of tumor establishment via BLI, mice were randomized into PBS control and treatment groups on Day 9 (**Fig. S2B**). Considering the inherent sensitivity of GBM8 cell line to TMZ, we chose two different TMZ doses in this model; the first being low dose (1 mg/kg) and the second being high dose (5 mg/kg). For the dose of EdU treatment, we used 200 mg/kg again as in U87 *in vivo* model, considering the fast clearance of the molecule. The combination groups featured combination of 1 mg/kg TMZ and 200 mg/kg and the combination of 5 mg/kg TMZ and 200 mg/kg, separately. As in the U87 *in vivo* model, 6 weeks of injections were administered with the M-F and MWF treatment regimens for EdU and TMZ, respectively. Unlike the U87 *in vivo* model, following the end of 6 weeks of injections, we wanted to investigate tumor recurrence and response to maintenance therapy in all treatment groups. Thus, we set a recurrence criteria per mouse where treatment was resumed when the tumor’s BLI signal increased to ≥10× its baseline (initial) value and a discontinuation criteria per mouse where reinjections were discontinued once the tumor’s BLI signal returned to and stabilized at the baseline level. The tumor growth was assessed via BLI in two phases: first phase during the administration of 6 weeks of injections (Day 7 to Day 50) and the second phase during the monitoring of tumor recurrence and response to as-needed maintenance therapy (Day 50 to study endpoint, Day 170).

### Assessment of tumor growth and recurrence during the first phase of treatment and the second phase of maintenance therapy in mice bearing orthotopic xenografts of GBM8

During the first phase, we observed suppression of tumor growth in all mice except the ones that received PBS and 1 mg/kg TMZ (**Fig. 5A, 5C, S2B, S2C, S3A-F**). During the second phase, when the initial treatment regimen had been discontinued, we started to observe recurrence in some mice in the 5 mg/kg TMZ, 200 mg/kg EdU, and 1 mg/kg TMZ + 200 mg/kg EdU groups and initiated maintenance therapy on recurrent mice on Days 59 and 63 (**Fig. 5B**, **S3B, S3C, S3F**). Recurrence occurred in ∼60% (4 out of 7) of mice in the 5 mg/kg TMZ group, 100% (6 out of 6) of mice in the 200 mg/kg EdU group, and ∼15% (1 out of 7) of mice in the 1 mg/kg TMZ + 200 mg/kg EdU group. In contrast, no recurrence was observed in any mice in the 5 mg/kg TMZ + 200 mg/kg EdU group (**Fig. 5B, S3D)**. According to the tumors’ BLI signals, none of the recurred mice in the 5 mg/kg TMZ group responded to maintenance therapy while 50% of the recurred mice in the 200 mg/kg EdU group responded to maintenance therapy and showed remission (**Fig. 5B, S3B, S3C**). One case of recurrence observed in the 1 mg/kg TMZ + 200 mg/kg EdU group also showed suppression of tumor growth and eventually remission upon receiving maintenance therapy (**Fig. 5B, 5C** and **S3F**; see **Fig. S2B-G** for extended BLI images). From these results, we concluded that maintenance therapy with single-agent EdU and combination of TMZ and EdU helped with suppressing the tumors while maintenance therapy with single-agent TMZ did not provide any benefit in suppressing the tumors.

**Figure 5:**
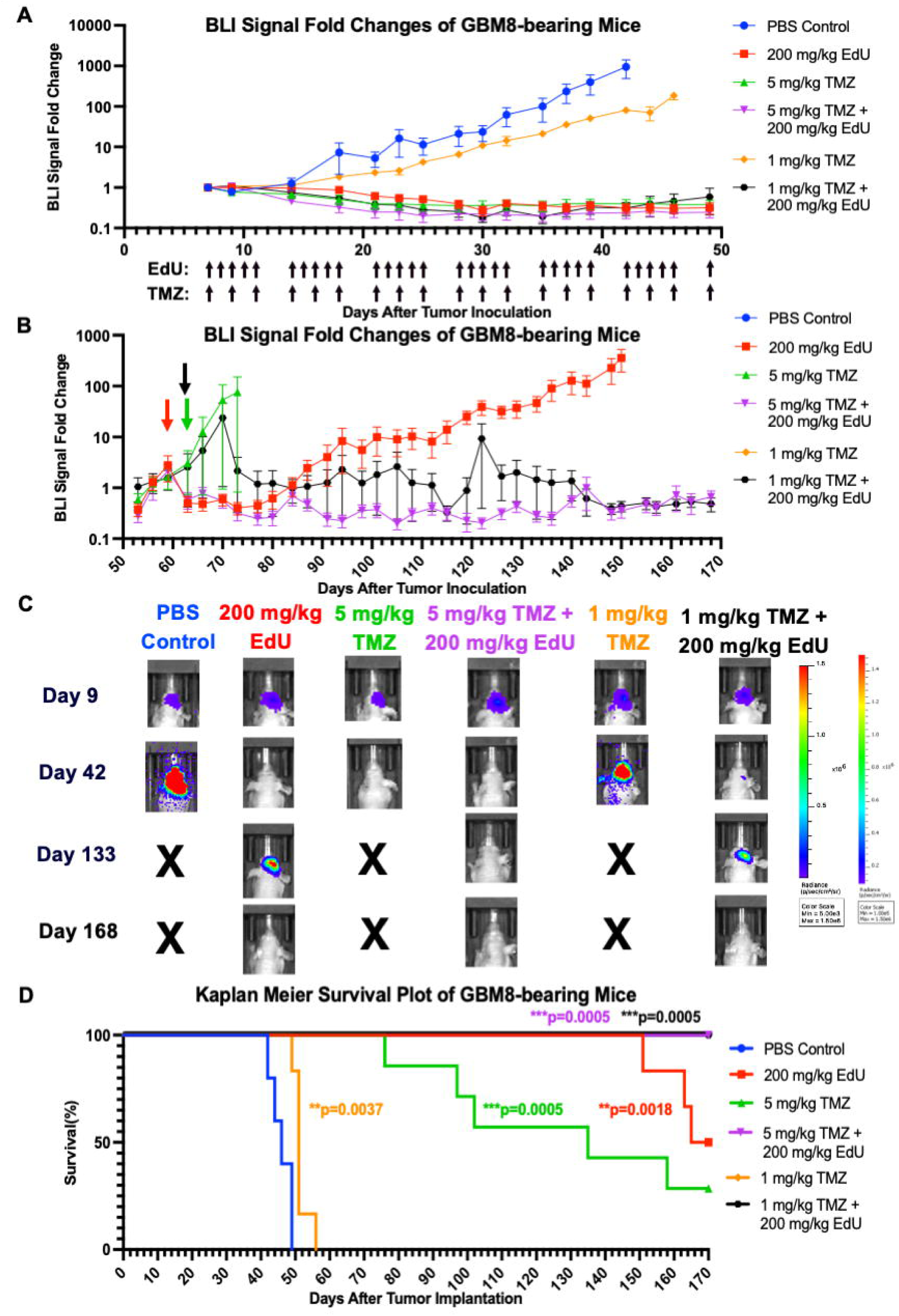
Testing the efficacy of TMZ+EdU in *in vivo* orthotopic GBM8 mouse model of GBM. **(A)** Bioluminescence imaging (BLI) signal fold changes of the tumors of mice bearing orthotopic xenografts of GBM8 and treated with either PBS (n=5), 200 mg/kg EdU (n=6), 5 mg/kg TMZ (n=7), 5 mg/kg TMZ + 200 mg/kg EdU (n=7), 1 mg/kg TMZ (n=6), or 1 mg/kg TMZ + 200 mg/kg EdU (n=7) during the first phase of treatment (day 0 to Day 50). Arrows under the x axis denote the days on which six weeks of EdU and TMZ injections were administered. **(B)** Bioluminescence imaging (BLI) signal fold change of the tumors of mice bearing orthotopic xenografts of GBM8 and treated with either PBS (n=5), 200 mg/kg EdU (n=6), 5 mg/kg TMZ (n=7), 5 mg/kg TMZ + 200 mg/kg EdU (n=7), 1 mg/kg TMZ (n=6), or 1 mg/kg TMZ + 200 mg/kg EdU (n=7) during the second phase of maintenance therapy (day 50 to Day 170). Arrows on the graph denote the day of the initiation of maintenance therapy in corresponding color-coded group. **(C)** Representative BLI images of mice bearing orthotopic xenografts of GBM8 and treated with either PBS, 200 mg/kg EdU, 5 mg/kg TMZ, 5 mg/kg TMZ + 200 mg/kg EdU, 1 mg/kg TMZ, or 1mg/kg TMZ + 200 mg/kg EdU on Days 9, 42, 133, and 168. **(D)** Kaplan Meier survival plot of mice bearing orthotopic xenografts of GBM8 and treated with either PBS, 200 mg/kg EdU, 5 mg/kg TMZ, 5 mg/kg TMZ + 200 mg/kg EdU, 1 mg/kg TMZ, or 1mg/kg TMZ + 200 mg/kg EdU (log-rank test).

### Monitoring survival in mice bearing orthotopic xenografts of GBM8

In addition to assessing tumor growth and recurrence, we also monitored survival throughout the study in the *in vivo* GBM8 model. At the study endpoint, single-agent treatments with 1 mg/kg TMZ, 5 mg/kg TMZ, and 200 mg/kg EdU extended the survival significantly by ∼10%, ∼200%, and ∼260%, respectively, compared to the PBS control group (p=0.0037, p=0.0005 and p=0.0011, respectively, **Fig. 5D**). Moreover, 50% of mice (3 out of 6 mice) in 200 mg/kg EdU group and ∼30% of mice (2 out of 7 mice) in 5 mg/kg TMZ group were alive by the study endpoint.

Importantly, all of the mice in 1 mg/kg TMZ + 200 mg/kg EdU and 5 mg/kg TMZ + 200 mg/kg combination groups were alive and tumor-free by the study endpoint, conferring a complete survival at the study endpoint in this stringent patient-derived model and highlighting the strong therapeutic potential of TMZ+EdU combination (**Fig. 5D**). We concluded from these results that the combination of TMZ and EdU conferred a synergistic effect in terms of both suppressing the tumor growth and extending the survival in mice bearing orthotopic xenografts of GBM8 cells. Combined with the results of the U87 *in vivo* model, these results supported our hypothesis that combining TMZ and EdU could yield a greater therapeutic effect than either of these drugs confers alone.

### Testing TMZ+EdU combination therapy in vitro in LN229 cell line and in mice bearing orthotopic xenografts of LN229

Upon observing these synergistic results with the GBM8 *in vivo* model, we wished to test this synergistic effect in a third orthotopic *in vivo* model using LN229 cell line. Before establishing the *in vivo* orthotopic model with this cell line, we performed cell survival assay to test the synergistic effect of TMZ+EdU *in vitro* by combining the IC50 value of EdU for LN229 (∼20 μM, as identified previously (25)) with various doses of TMZ. As in the other two GBM cell lines, the TMZ+EdU combination was more efficient than either drug alone at higher concentrations of TMZ (**Fig S4A**). These synergistic results led us to test the synergism of TMZ+EdU combination in the *in vivo* orthotopic model with LN229 cell line.

Accordingly, we created the *in vivo* orthotopic model by inoculating LN229 cells intracranially into the brain parenchyma of female athymic nude mice. Upon confirming the establishment of tumors via BLI, we stratified the mice into PBS control and five treatment groups on Day 4 (**Fig. S4B**). We chose the same treatment groups and treatment regimen with 6 weeks of injections as in the GBM8 *in vivo* model. By the third week of injections, mice in PBS control group started to reach humane end point while the tumors’ BLI signals in all the other groups, except 200 mg/kg EdU group, decreased below the baseline level, indicating suppression of tumor growth (**Fig. 6A, 6B, S4C**). On the other hand, 200 mg/kg EdU treatment was only able to stabilize the tumors’ BLI signals at ∼3 times the baseline level signal. By the completion of 6 weeks of injections on Day 46, all the mice in the PBS control group reached humane endpoint and the tumors’ BLI signals in 200 mg/kg EdU groups started to increase. However, the tumors’ BLI signals in TMZ single-agent and TMZ+EdU combination groups decreased further and stabilized ∼0.01 times the baseline levels (**Fig. 6A**). Around a month after the completion of injections by Day 70, mice treated with 200 mg/kg EdU started to reach humane endpoint while tumors’ BLI signals in combination or single-agent TMZ groups remained lower than the baseline level signal (**Fig. 6A, S4D-F**). Additionally, around Day 80 in 25% (2 out of 8 mice) of mice in 1 m/kg TMZ group and on Day 85 in ∼12% (1 out of 8 mice) of mice in 5 mg/kg TMZ group, we started to observe some recurrence (**Fig. 6A, 6B, S4G**). However, this recurrence was not observed in either of the combination groups. For survival, treatment with 200 mg/kg EdU extended the median survival significantly by ∼110% compared to the PBS control group (p=0.0002, **Fig. 6C**). At conclusion of the study, none of the mice in 5 mg/kg TMZ + 200 mg/kg EdU group reached the humane endpoint, resulting in a striking and significant increase in median survival compared to both PBS control and single-agent EdU treatment groups (p<0.0001 and p=0.0008, respectively, **Fig.6C**). Similarly, 1 mg/kg TMZ + 200 mg/kg EdU group extended the survival significantly compared to both PBS control and single-agent EdU group (p<0.0001 and p=0.0018, respectively, **Fig.6C**). In 1 mg/kg TMZ + 200 mg/kg EdU group, only one mouse reached humane endpoint due to orbital infection, although its tumor had way lower BLI signal compared to the baseline level, corresponding to ∼88% survival at the study endpoint. However, different from our results in U87 and GBM8 *in vivo* models, in the LN229 *in vivo* model, we observed that combination of TMZ and EdU performed as well as the single-agent TMZ groups, leading to an additive effect in suppressing the tumor growth and prolonging the median survival (**Fig. 6C**). Collectively, the results from our *in vivo* orthotopic mouse models of GBM established with U87, GBM8 and LN229 cells provided preclinical evidence that TMZ+EdU combination could elicit a synergistic to partly additive effect in GBM, supporting the progression of this combination to early-phase clinical testing.

**Figure 6:**
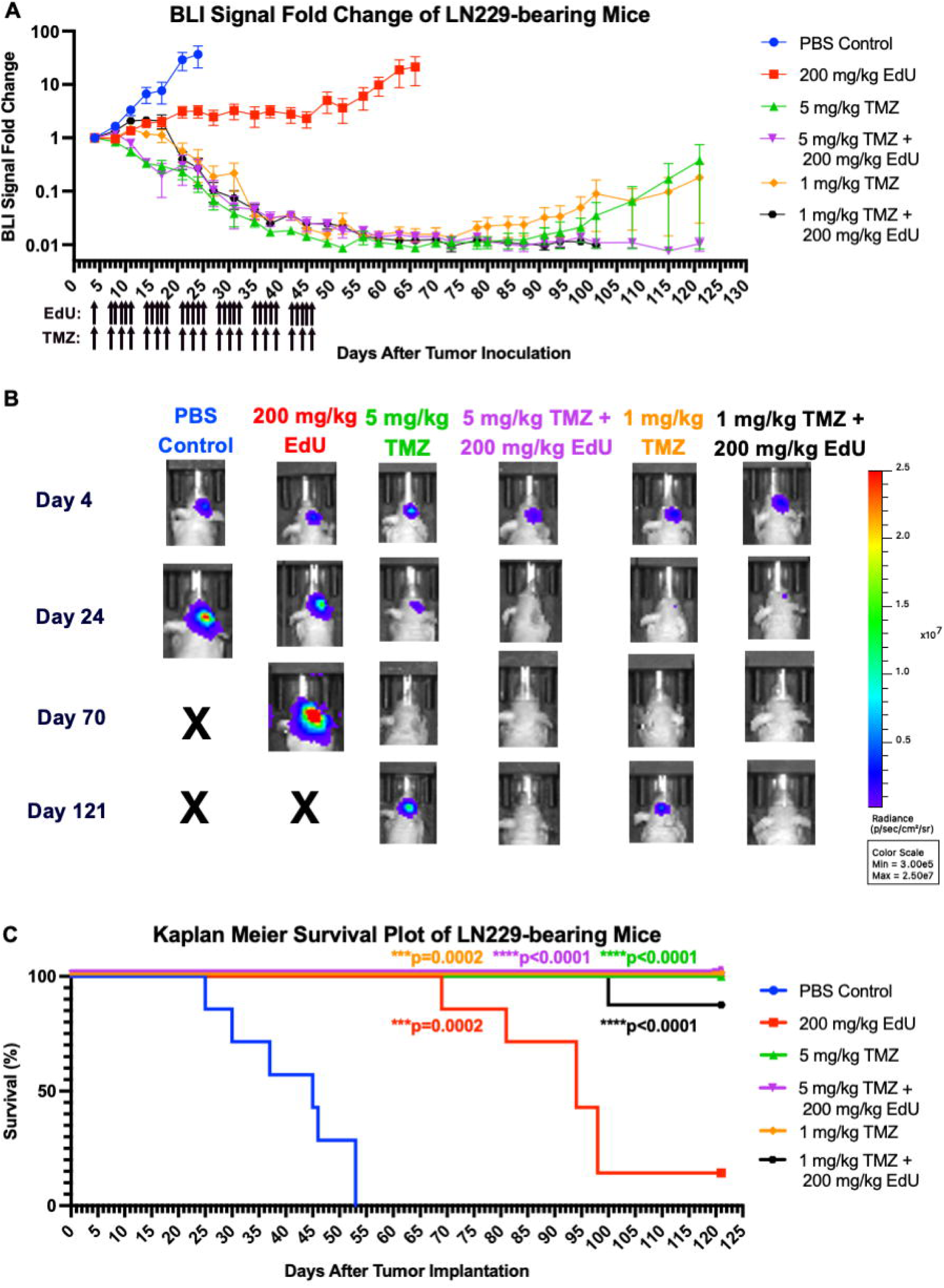
Testing the efficacy of TMZ+EdU in *in vivo* orthotopic LN229 mouse model of GBM. **(A)** Bioluminescence imaging (BLI) signal fold changes of the tumors of mice bearing orthotopic xenografts of LN229 and treated with either PBS (n=7), 200 mg/kg EdU (n=7), 5 mg/kg TMZ (n=8), 5 mg/kg TMZ + 200 mg/kg EdU (n=8), 1 mg/kg TMZ (n=7), or 1 mg/kg TMZ + 200 mg/kg EdU (n=8). Arrows under the x axis denote the days on which six weeks of EdU and TMZ injections were administered. **(B)** Representative BLI images of mice treated with either PBS, 200 mg/kg EdU, 5 mg/kg TMZ, 5 mg/kg TMZ + 200 mg/kg EdU, 1 mg/kg TMZ, or 1 mg/kg TMZ + 200 mg/kg EdU on Days 4, 24, 70, and 98. **(C)** Kaplan Meier survival plot of mice bearing orthotopic xenografts of LN229 and treated with either PBS, 200 mg/kg EdU, 5 mg/kg TMZ, 5 mg/kg TMZ + 200 mg/kg EdU, 1 mg/kg TMZ, or 1 mg/kg TMZ + 200 mg/kg EdU (log-rank test).

### Toxicity profiling of mice treated with EdU, TMZ and their combination in *in vivo* orthotopic mouse model of U87

To assess treatment-associated toxicity, we collected peripheral organs (intestine, kidney, spleen, and liver) from mice bearing orthotopic xenografts of U87 at the humane endpoint. Organs were fixed, processed, paraffin-embedded, sectioned, and stained with hematoxylin and eosin (H&E). Slides were histopathologically evaluated and scored for organ-specific signs of toxicity (**Fig. S5**). In the small intestine of U87 tumor-bearing mice, minimal to mild crypt epithelial necrosis/apoptosis was observed in 8 of 10 animals treated with 200 mg/kg EdU. Regenerative activity, evidenced by increased mitotic figures, was also present in some animals. In mice treated with 5 mg/kg TMZ + 200 mg/kg EdU, no overt crypt necrosis/apoptosis was seen, but increased mitotic figures were detected in 2 of 5 animals, suggesting earlier crypt injury followed by regeneration. These mice survived ∼30 days longer than EdU-only animals, supporting the interpretation that the intestinal epithelium had recovered from prior EdU-related injury. No crypt changes were observed in mice treated with 5 mg/kg TMZ or PBS. The mild severity and evident reversibility of the crypt epithelial injury indicate a non-adverse effect. In the kidney, minimal multifocal cortical tubular epithelial degeneration/regeneration was noted in 2 of 10 EdU-treated animals and 1 of 5 animals receiving the TMZ+EdU combination, but not in TMZ-only or PBS-treated mice. As this lesion is a common background finding and may occur with general debilitation, its low incidence and severity suggest a non-adverse, possibly incidental effect. In the spleen, increased extramedullary hematopoiesis (EMH) was observed in 7 of 10 mice treated with EdU and 4 of 5 mice treated with TMZ+EdU, but not in TMZ-only animals and only minimally in 1 of 7 PBS controls. This finding is considered EdU-related and likely reflects bone marrow suppression. Slight decreases in splenic lymphocyte cellularity were also seen in EdU- and combination-treated mice; however, similar changes occurred in PBS controls, suggesting a stress-related rather than treatment-specific effect. In the liver, no hepatocellular injury was detected. Minor group differences in hepatocellular vacuolation were consistent with expected diurnal variation in glycogen content (hepatocytes from animals euthanized earlier in the day typically exhibit greater glycogen-associated vacuolation than those euthanized later) (35–38). Overall, these observations demonstrate that EdU, alone or in combination with TMZ, induced only mild and reversible histopathological changes in peripheral organs. This conclusion is also supported by the recovery of body weight after treatment in mice bearing U87, GBM8, and LN229 tumors (**Fig. S6A-C**).

In addition to these toxicity profiling of the peripheral organs, we also wanted to test the effect of the treatment with EdU, TMZ and their combination on complete blood counts. To this end, we collected blood samples from mice bearing LN229 tumors on three different days: two of them during the 6 weeks of injections (Day 16 and Day 30) and one after the completion of 6 weeks of injections (Day 58). According to the results of complete blood counts, there were only minor and reversible differences between the counts of White Blood Cells (WBCs), Lymphocytes (LYMPH), Neutrophils (NEUT), Basophils (BASO), Eosinophils (EO), and Red Blood Cells (RBC) values in PBS control group on Day 16 and in all the other groups. Additionally, for all the counts of these different cells, the values in all groups were mainly at or above the normal lower limit values by Day 58 (**Fig. S7A-F**). These results, together with the results on the histopathological and body weight changes, further confirmed that EdU, alone or in combination with TMZ, induced mild, transient, and reversible toxicity.

### Testing TMZ+EdU combination therapy against living, passage-zero GBM patient tumor tissues *ex vivo*

Upon collecting these promising *in vivo* results, we proceeded to test the efficacy of TMZ+EdU on living, passage-zero GBM patient tissues seeded atop organotypic brain slice cultures (25,39,40). In our previous investigation of EdU as a potential therapeutic for GBM, we observed differential response across all patient tumors tested with monotherapy EdU or TMZ (25). In these new experiments, we observed a similar trend. We exposed four IDH-wild-type WHO-classified uncultured patient GBM tissues to single-agent EdU or TMZ as well as a combination of both compounds. 3 out of 4 of these tissue samples (PTS125, PTS152, and PTS153) were from patients who clinically showed rapid tumor progression after receiving TMZ.

During treatment, TMZ and/or EdU were added under the transwell membranes supporting each tumor-bearing OBSC and allowed to diffuse through the OBSCs and into the tumor cells. All tumors were treated with EdU and/or TMZ at doses of 0, 100, 250, and 1000µM. Combination therapy of [TMZ + EdU] was given at doses of [0, 0], [100, 100], [250, 250] and [500 µM, 500 µM]. Among the tissues tested, we observed several doses where EdU alone or in combination with TMZ performed better than TMZ alone (**Fig. 7A-D**). Using the Zero Interaction Potency (ZIP) Synergy score (41,42) that measures the difference between observed effects of a combinatorial regimen and expected effects if individual drugs are not interactive, we found that the effect of the combinatorial therapy ranged from additive (0 ˂ ZIP < 10 in PT158, PT125, PT152) to synergistic (ZIP > 10 in PT153) (**Fig. 7E**).

**Figure 7:**
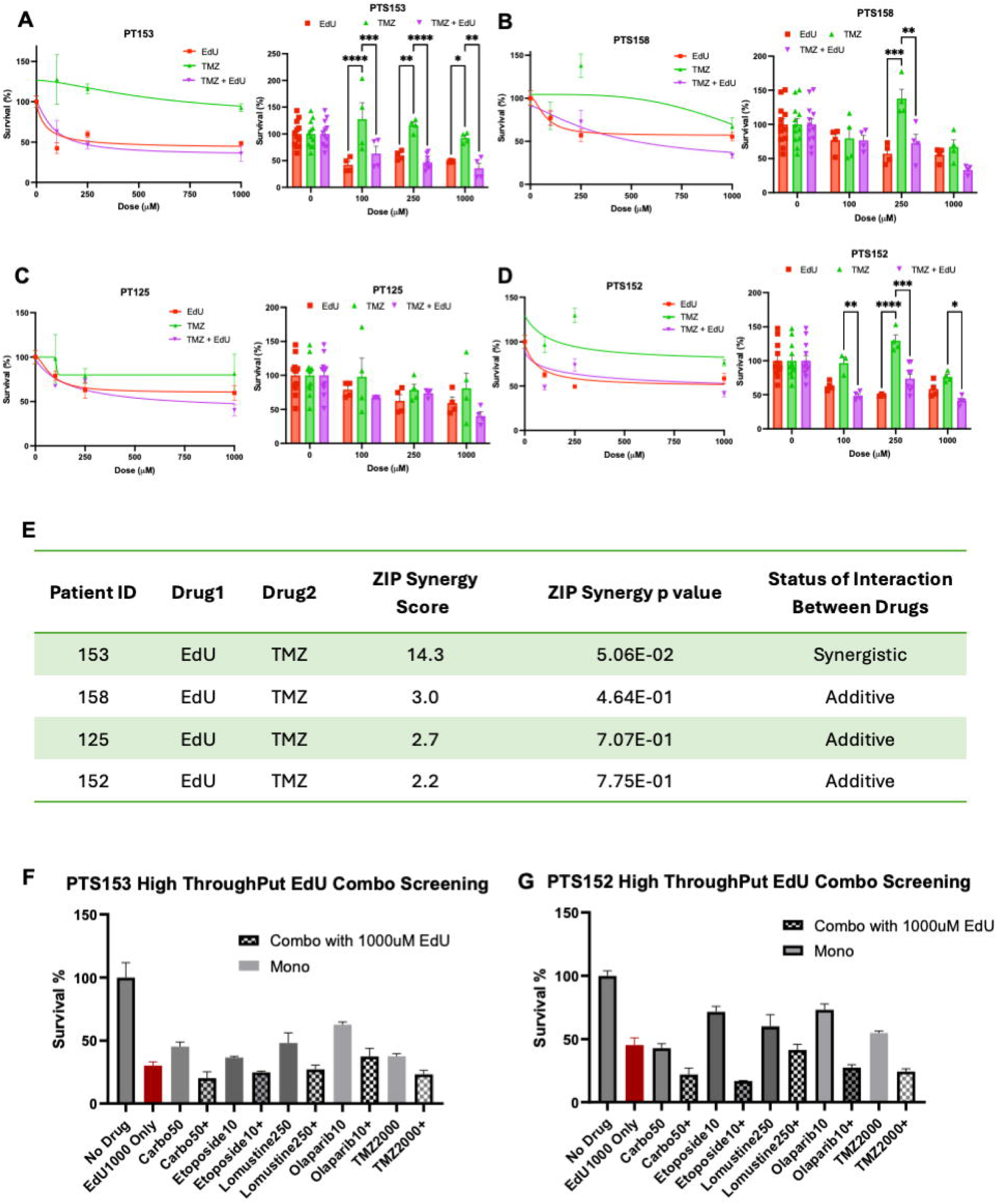
Testing the efficacy of TMZ+EdU in living, uncultured GBM patient tissues seeded on OBSC. Dose-response curves and grouped dose-response bar graphs of GBM patient tissues PT153 **(A)**, PT158 **(B)**, PT125 **(C)**, and PT152 **(D**) treated with single-agent EdU, TMZ or TMZ+EdU (Two-way ANOVA, ****p<0.0001, ***p<0.001). **(E**) Zero Interaction Potency (ZIP) Synergy Score values and ZIP p values for TMZ+EdU combination tested on four GBM patient tissues. High-throughput combo screening of PT153 **(F)** and PT153 **(G)** with 1000 μM EdU in combination with carboplatin (50 μM), etoposide (10 μM), lomustine (250 μM), Olaparib (10 μM), and TMZ (2000 μM).

GBM tumors have significant intra- and inter-tumoral heterogeneity; therapies that are effective for some patients may not be effective for others, and vice versa. We therefore tested EdU against two passage-zero patient GBM tissues in combination with a range of other FDA-approved chemotherapies. We chose carboplatin and lomustine because of their cell cycle-independent anti-tumor activity to complement EdU’s hypothesized activity during the S-phase (43,44), and also because lomustine is frequently used as a control arm in trials and sometimes considered the standard of care after GBM progression (43,45). We chose etoposide because its mechanism of action through the stabilization of the topoisomerase II-DNA cleavage complex is very different from EdU’s and may elicit different downstream compensatory mechanisms. It has also been well-tolerated in heavily pretreated patients with refractory gliomas (46,47). We also chose the poly (ADP-ribose) polymerase-1 inhibitor (PARPi) Olaparib because it is under investigation as a radio and chemosensitizer (48,49). As such, it might potentiate EdU’s ability to target glioma stem cells (GSCs) which have upregulated PARP that promotes base excision repair and the repair of single-strand breaks in treated cancer cells (50).

To assess these exploratory combinatorial regimens, we first used doses previously determined to be close to the ED50 values of each single agent. We then combined these drugs with 1000 µM EdU. Across all drugs tested in combination, we observed more killing in the combination groups than when the drug was used alone (**Fig. 7F, 7G**). Notably, at the same doses, each patient tissue responded differently to each drug. For instance, in PTS152, there was a 2.9-fold increase in tumor killing when EdU was added to etoposide, but just a 1.2 fold increase in PTS153, (**Fig. 7F, 7G**), further highlighting the need for more personalized and patient-centric models that capture patient-to-patient heterogeneity.

## DISCUSSION

As the most common primary malignancy of the central nervous system, GBM carries an extremely poor prognosis, with only ∼7% of patients surviving five years after diagnosis (51–53). Therefore, there is an urgent need to develop new treatment modalities for GBM. Current efforts focus on investigating the therapeutic potential of integrating different drugs into combinatorial treatments with TMZ, the first-line chemotherapeutic for GBM. However, despite trials and preclinical studies with agents such as lomustine (54), bevacizumab (55), and PARP inhibitors (56,57), none of these combinations have received FDA approval to date.

In this work, we demonstrated in (i) GBM cell lines *in vitro*, (ii) orthotopic mouse models of GBM *in vivo*, and (iii) living, uncultured GBM patient tissues on OBSC that EdU shows a synergistic effect in GBM when combined with TMZ. Moreover, for the first time, we characterized the pharmacokinetic features of EdU and confirmed the targeted localization of EdU into GBM tumor cells *in vivo* in both single-agent EdU-treated and combination-treated mice.

Our results on pharmacokinetic characteristics of EdU *in vivo* highlighted the fast clearance of EdU with limited penetrance to the blood brain barrier and suggested that there is room for improvement to modulate the bioavailability of EdU via formulation. Formulating EdU would increase the bioavailability and in turn the penetrance of EdU to blood brain barrier, potentiating the therapeutic effect we observed in combination with TMZ.

In our U87 *in vivo* model, we observed only a partial increase in survival with single-agent TMZ and EdU while neither was able to suppress the tumor growth. However, combination of TMZ and EdU showed a truly synergistic effect in this model by both suppressing the tumor growth and extending the survival drastically. On the other hand, in our GBM8 *in vivo* model, single-agent EdU and TMZ (5 mg/kg) treatments conferred considerable survival benefit compared to PBS control group. Nevertheless, we also observed recurrence in these single-agent treatment groups with partial to no response to maintenance therapy. For instance, we observed recurrence and no response to maintenance therapy in ∼60% of the mice treated with 5 mg/kg TMZ, reflecting the acquired resistance following an initial positive response to TMZ treatment. Notably, this TMZ resistance was overcome in mice treated with 5 mg/kg TMZ + 200 mg/kg EdU since no recurrence was observed in these mice by the study endpoint. These results point out the power of TMZ+EdU combination in targeting the TMZ-resistant tumors which is a prevalent occurrence in the clinic.

Surprisingly, in our third *in vivo* model with LN229, we observed that single-agent TMZ treatments initially performed as well as the combination treatment groups, indicating an additive effect in this model. However, both single-agent TMZ groups exhibited recurrence following the completion of six weeks of injections. Combination groups in this LN229 model, on the other hand, were still able to avoid this recurrence, similar to the results in GBM8 model while the survival benefit these combination treatments conferred was not as drastic as in GBM8 model. The difference among these different *in vivo* models of GBM could be attributed to the heterogeneous nature of this disease and the differences among the genetic alterations harbored by these GBM cell lines. Additionally, further *in vivo* studies in these and other models featuring various doses of EdU in combination with TMZ could identify lower doses of EdU that could still show synergistic effect.

Furthermore, on our OBSC platform, TMZ+EdU acted in an additive or synergistic manner against all four passage-zero patient tumor tissues. Notably, these findings demonstrate that the positive interactions between EdU and TMZ extend beyond *in vitro* and animal models to actual heterogeneous human tumors. The more muted responses measured here reflect what one may expect to observe in the clinical setting, where tumors with heterogeneity in proliferation rate and mutational drivers are typically not responsive to TMZ alone and respond incrementally to combination therapies. The heterogeneity of patient-to-patient response also highlights the need for more personalized and patient-centric models in drug development. Our OBSC platform may therefore serve as a functional correlative biomarker to identify patients most likely to benefit from TMZ+EdU or other combinatorial regimens. Expanding OBSC testing of EdU to a larger, molecularly diverse cohort could refine the estimated prevalence of synergy, enable biomarker discovery, and further advance functional precision medicine efforts in GBM.

### New therapeutic avenues to pursue: EdU’s potential in combination with various other chemotherapeutics

Our results on combining EdU with various other chemotherapeutics against GBM patient tumor tissues revealed the transformative role EdU could play in alternative treatment regimens for GBM. By targeting complementary mechanisms of action in these combinations, we found patient-specific sensitivities to different combination therapies. Future studies further testing the mechanism and therapeutic potential of these combinatorial regimens will shed light on the clinical utility of each in GBM. Introducing such personalized combination regimens into clinical GBM treatment could provide much-needed alternatives for patients with GBM.

## MATERIALS & METHODS

### Cell Lines

U87 (purchased from ATCC in 2012, catalog #: HTB-14, RRID: CVCL_0022) and LN229 (purchased from ATCC in 2012, catalog #: CRL-2611, RRID: CVCL_0393) cells were cultured in 1X DMEM supplemented with 10% fetal bovine serum (FBS) and 1% penicillin-streptomycin in T75 flasks. Patient-derived GBM8 cells (a kind gift from gift from H. Wakimoto, Massachusetts General Hospital) (29,30) were cultured in neurobasal-A medium (Gibco, catalog #: 12349015) supplemented with 1 mg heparin (STEMCELL Technologies, catalog #: 07980), 10 μg FGF (Shenandoah Biotechnology, catalog #: 100-146), 10 mL B27 supplement (Life Technologies Inc., catalog #: A3582801), 7.5 mL L-glutamine (Life Technologies Inc., catalog #: 10888022), 10 μg EGF (Shenandoah Biotechnology, catalog #: 100-26-100ug), 2.5 mL N2 supplement (Life Technologies Inc., catalog #: 17502048), and 2.5 mL Antibiotic-Antimycotic (100X) (Thermo Fisher Scientific, catalog #: 15240062) in T75 flasks.

### Lentiviral vectors

The following lentiviral vector (LV) was used in this study: mCherry protein fused to firefly luciferase (LV–mCh-FL). This LV also encodes a puromycin resistance gene. U87, GBM8 and LN229 cell lines were transduced with this mCherry-FLuc Lentivirus. Following transduction, mCherry expression was confirmed and cells expressing mCherry were selected using puromycin treatment.

### *In vitro* survival assay via BLI

The cells were counted using a Countess Cell Counter (Thermo Fisher Scientific, catalog #: AMQAX1000) and seeded in triplicate with a 2500 cells/well density in black bottom 96 well plate. Following 24 h incubation, the cells were treated with increasing doses of single agent TMZ (Sigma-Aldrich, catalog #: T2577-100MG), single agent EdU (Sigma-Aldrich, catalog #: 900584-500MG). For combination, IC50 value of the cell line for EdU was combined with increasing doses of TMZ. At the end of 72 h treatment, the cell viability was measured via BLI using an AMI optical imaging system. The data was analyzed using Aura software, Microsoft Excel, and GraphPad Prism to plot dose-response curves (‘Non-linear Regression (curve fit)’ option used in GraphPad Prism).

### Assessing synergism between TMZ and EdU *in vitro*

The cell viability data obtained from 3 biological replicates of *in vitro* survival assay via BLI of U87, LN229 and GBM8 cell lines for each single dose of single agent EdU, TMZ or their combination were fed in CompuSyn software as Fa (fraction affected). The CompuSyn software then generated “Logarithmic Combination Index Plot” for drug combinations for each cell line where log of Combination Index (CI) is plotted against the Fa (58–60). In this plot, the log(CI)<0, log(CI)=0 and log(CI)>0 show synergism, additive effect and antagonism between EdU and TMZ, respectively.

### Establishment of *in vivo* orthotopic xenograft mouse models of human GBM

All animal experiments were conducted in accordance with institutional guidelines and approved by the Institutional Animal Care and Use Committee (IACUC) at the University of North Carolina at Chapel Hill (UNC-CH) under UNC IACUC protocol 25-016. Female athymic nude mice (in-house bred) were housed under sterile conditions in an AAALAC-accredited animal facility at UNC-CH.

U87, LN229, and GBM8 glioma cells expressing luciferase were cultured and prepared for intracranial (i.c.) inoculation into the brains of athymic female nude mice. U87 and LN229 cells were detached using 0.05% Trypsin-EDTA (1X; Gibco, catalog #25300-054) and incubated for 5 minutes. GBM8 cells were detached using 1X Accutase (STEMCELL Technologies, catalog #: 07920) and incubated for 5 minutes, with declumping by pipetting at 2.5 minutes. After detachment, cells were counted using a Countess II Cell Counter, and suspensions were prepared in PBS at final concentrations of 3.3 × 10⁴ cells/μL (U87), 5 × 10⁴ cells/μL (GBM8), and 8.3 × 10⁴ cells/μL (LN229). For intracranial inoculation, mice were anesthetized with isoflurane (2.0–2.5% for induction; 2.5–3.0% for maintenance) and placed in a stereotactic apparatus following confirmation of adequate anesthesia via a negative toe-pinch reflex. After surgical site preparation, a midline scalp incision was made (<10 mm) and a ∼1 mm burr hole was drilled to facilitate needle insertion for intracranial injection. Each mouse was injected with a 3 μL volume of the corresponding cell suspension of U87, GBM8 or LN229 cells using the following coordinates relative to bregma: mediolateral (ML) = 2.5 mm, anteroposterior (AP) = 0.5 mm, dorsoventral (DV) = 2 mm. The total cells inoculated were 100,000 cells per mouse for U87 (n=40), 250,000 cells per mouse for LN229 (n=50), and 150,000 cells per mouse for GBM8 (n=42). The wounds were closed with 3M Vetbond™ tissue adhesive. Mice were monitored during and post-surgery and received meloxicam once daily for 48 hours postoperatively as established in the approved animal protocol.

### Monitoring tumor growth via BLI in *in vivo* orthotopic xenograft mouse models of human GBM

Four to nine days post-implantation of U87, GBM8 or LN229 cells into the brains of mice, mice were injected with IVISbrite D-Luciferin Potassium Salt Bioluminescent Substrate (1g, XenoLight) (Revvity, catalog #: 122799-5) at 150 mg/kg, anesthetized with isoflurane and imaged via bioluminescence imaging (BLI) with IVIS Spectrum optical imaging system. The BLI signals recorded were analyzed using the LivingImage software and mice were stratified into treatment groups based on tumor burden as assessed by BLI. The monitoring of tumor growth in all of these *in vivo* models continued throughout the study durations at a frequency of 2-3 times a week.

### Administration of EdU, Temozolomide (TMZ), and their combination, monitoring of tumor growth and evaluation of therapeutic response in *in vivo* orthotopic xenograft mouse models of human GBM

For the U87 and LN229 models, mice received a total of 6 weeks of injections. The treatment regimen for EdU was 200 mg/kg EdU once every weekday (M-F) and for TMZ either 1 mg/kg or 5 mg/kg TMZ every other weekday (MWF). For the combination group, the mice received TMZ injections in the morning and EdU injections in the afternoon with the same regimens. The tumor growth was monitored via bioluminescence imaging (BLI) of the mice using IVIS Spectrum device through intraperitoneally injecting luciferin. The mice started to receive treatment with EdU, TMZ and their combination 6 days and 4 days post tumor cell inoculation for U87 model and LN229 model, respectively. For the GBM8 model, the mice started to receive 6 weeks of injections with the abovementioned regimens for EdU, TMZ and their combination 7 days post tumor cell inoculation and the tumor growth was monitored similarly. Different from the other models, the injections restarted once a mouse’s tumor recurred in GBM8 model. The recurrence criterion was recording a BLI signal that is 10 times higher than the initial signal recorded 7 days post tumor cell inoculation. The injections continued until a consistently low BLI signal (4 BLI readings showing either the same initial BLI signal or lower). The injections were discontinued once this stopping criterion was met. Tumor growth was continuously monitored by BLI throughout the duration of the studies in all models. In the U87 and LN229 models, mice underwent a 6-week treatment regimen. EdU was administered at 200 mg/kg once daily on weekdays (Monday– Friday), while TMZ was given at either 1 mg/kg or 5 mg/kg every other weekday (Monday, Wednesday, Friday). For combination treatment, mice received TMZ injections in the morning and EdU in the afternoon, following the same individual dosing schedules. Tumor growth was monitored via bioluminescence imaging (BLI) using the IVIS Spectrum system, following intraperitoneal injection of luciferin. Treatment began 6 days post–tumor cell inoculation for the U87 model and 4 days post-inoculation for the LN229 model. In the GBM8 model, treatment also began 7 days after tumor cell inoculation and followed the same 6-week regimen. Tumor progression was monitored via BLI, as in the other models. Unique to GBM8, treatment resumed upon tumor recurrence, defined as a BLI signal ≥10x the baseline signal recorded on day 7 post-inoculation. Injections continued until the BLI signal returned to and remained at baseline or lower for four consecutive readings, at which point treatment was discontinued. Tumor growth was continuously monitored by BLI throughout the duration of the studies in all models, along with survival tracking. Mice were also observed daily or every other day for signs of neurological symptoms, ≥20% body weight loss, or decline in body condition score. Animals exhibiting any of these signs were euthanized. Survival analysis was performed using the log-rank (Mantel–Cox) test in GraphPad Prism.

### Blood collection from mice bearing orthotopic xenografts of LN229 cells for complete blood counts

Blood samples were obtained via a small venous nick on the mouse tail using a 22-gauge needle. This procedure was performed on mice bearing orthotopic xenografts of LN229 cells and receiving different treatment regimens and complete blood counts were assessed on days 16, 30, and 58 following tumor implantations. Blood collection volumes adhered strictly to the approved animal use protocol. A maximum of three sampling sessions were conducted at bi-weekly intervals, with each individual collection limited to approximately 30-50 µL. Complete blood counts (CBCs) were performed on whole blood using an IDEXX ProCyte Dx™ Hematology Analyzer. Parameters measured included indices of leukocyte, erythrocyte, and platelet lineages.

### Processing, embedding, sectioning, staining and imaging of tissues from peripheral organs of mice bearing orthotopic xenografts of U87 and GBM8 Cells

Formalin-fixed tissues were processed in a Leica Pegasus tissue processer, embedded in paraffin (Leica Paraplast) using a HistoCore Arcadia H. Tissues were then sectioned on a microtome (Leica, catalog #: RM2255) at 5 microns onto positively charged slides (Fisher Scientific, catalog #: 12-550-15). For brains, the tissues were trimmed and sectioned around the site of tumor cell inoculation in the right hemisphere at coordinates of 0.2 mm anterior-posterior (AP), 2.0 mm mediolateral (ML), and 3.5 mm dorsoventral (DV).

### H&E staining and imaging

Prior to H&E staining, slides were air dried overnight and baked for 30 minutes at 60 degrees Celsius. H&E stains were performed using the Autostainer XL from Leica Biosystems. The slides were deparaffinized in graded ethanol, then stained with Hematoxylin (Epredia, catalog #: 7211) for 2 mins and Eosin -Y (Epredia, catalog #: 71311) for 1 min with respective washes in between. Clarifier 2 (catalog #: 7402) and Bluing (catalog #: 7111) solutions from Epredia were used to differentiate the reaction. After staining, slides were then dehydrated in graded ethanol, ending in xylene, and coverslipped with Surgipath Micromount (Leica, catalog #: 3801731). Stained slides were digitally imaged in the Aperio AT2 (Leica Microsystems) using 20x objective.

### EdU and DAPI staining and imaging

The slides with brain tissues at the site of tumor inoculation were deparaffinized in graded ethanol. They were then washed once with 1X Dulbecco’s PBS (Gibco, catalog #: 14190144) for 10 minutes and once with 0.1% Triton X-100 (Sigma-Aldrich, catalog #: T9284-500ML) in PBS for 10 minutes in Coplin jars at room temperature. The slides were then permeabilized, blocked, washed, and stained for EdU with Click-iT Plus Alexa Fluor picolyl azide toolkit (Thermo Fisher Scientific, catalog #: C10641) and for DAPI with 1 μg/ mL Hoechst 33258, pentahydrate (bis-benzimide) (Thermo Fisher Scientific, catalog #: H3569) as previously described (25). Upon completion of these stainings, the slides were mounted with ProLong Gold Antifade Mountant (Thermo Fisher Scientific, catalog #: P36934) and cured overnight.

The next day, slides were digitalized using the Aperio ScanScope FL (Aperio Technologies Inc). The digital images were captured in each channel by 20x objective (0.468 μm/pixel resolution) using line-scan camera technology (U.S. Patent 6,711,283). The adjacent 1 mm stripes captured across the entire slide were aligned into a contiguous digital image by an image composer. Images were archived in PSC’s eSlide Manger database (Leica Biosystems).

### OBSC generation

All animal work related to OBSC generation was approved by the Institutional Animal Care and Use Committee at the University of North Carolina at Chapel Hill under protocol 22-171. P8 Sprague-Dawley rat pups were used for all OBSC preparation. Animals were housed with one mother and no more than ten pups. All animals were euthanized prior to weaning. OBSCs were generated as previously described. In brief, 300um coronal sections of rat brains were generated with a VT1000S vibratome and suspended on a cell culture insert immersed in 1ml of brain slice media consisting of “Neurobasal-A medium supplemented with 10% heat-inactivated pig serum, 5% heat-inactivated rat serum, 1 mM L-glutamine, 10 mM KCl, 10 mM HEPES, 1 mM sodium pyruvate and 100 U/mL penicillin-streptomycin.” (39,40) The slices were then incubated at 37°C, 5% CO2 and 95% humidity.

### Patient tissue identification, procurement, and processing

All brain tumor specimens were collected at University of North Carolina Hospitals. A total of four patients with primary or recurrent IDH-wild-type WHO-classified GBM diagnosis were included in the present study. Appropriate written informed consent by parent/guardian and assent as applicable was obtained under protocols where ethical approval was given by the Institutional Review Board (IRB) in the Office of Human Research Ethics at the University of North Carolina at Chapel Hill. All patients consented to non-interventional IRB Protocol #23-0834 (LCCC2212; ClinicalTrials.gov NCT05978557), which began on 27/07/2023; estimated end date 01/03/2028. The laboratory study team had no access to identifying patient information; de-identified or coded diagnostic data describing tumor type and mutational status was provided by a Study Coordinator or Honest Broker.

Upon standard of care surgery, leftover tumor tissues not necessary for diagnosis were deidentified and transferred to the laboratory on wet ice. The tissue samples were immediately washed with PBS and red blood cells (RBC) were lysed with an RBC lysis buffer. The tissue samples were then mechanically dissociated with scalpels and cryopreserved at 150 mg of tissue per ml of CryoStor CS10.

### Patient tissue processing and drug killing

Originally cryopreserved patient tumor tissue samples were processed for drug dosing according to established protocols at the Screening Live Cancer Explants Core Facility at UNC (25,39). In brief, tissue samples were retrieved from liquid nitrogen storage, rapidly thawed, washed with PBS and passed through a 100um strainer to approximate a single-cell suspension. The tissue was then exposed to a LV–mCh-FL lentivirus to express both fluorescent mCherry and bioluminescent firefly luciferase reporters. On Day 0, 1 mg of tissue was seeded atop each hemisphere of brain slices suspended in a cell culture insert. On D1, corresponding doses of EdU, TMZ, carboplation (Sigma Aldrich, catalog #: C2538-100MG), lomustine (Selleck Chemicals, catalog #: S1840), Olaparib (Selleck Chemicals, catalog #: S1060), or etoposide (Sigma Aldrich, E1383-100MG) diluted in 1 ml of brain slice media were added underneath the inserts. The treated seeded tumors were then incubated for 3 days and bioluminescence readouts were obtained from an AMI optical imaging system.

### Synergy score calculation

All doses of drugs tested were exported into Synergy finder https://synergyfinder.org/ and synergy scores were automatically calculated. Synergy scores less than -10, between -10 and +10, and greater than +10 indicate antagonistic, additive, or synergistic interactions respectively.

### Pharmacokinetic Study

Pharmacokinetic (PK) analyses were conducted in healthy 12-week-old female Nu/Nu mice. Each animal received a single intraperitoneal (IP) dose of EdU (200 mg/kg). Blood and brain samples were collected at 15 min, 30 min, 1 h, 2 h, and 4 h after the dose (n = 5 per time point) and at 6 h (n = 3) under isoflurane anesthesia. Approximately 0.8–1.0 mL of blood was obtained via cardiac puncture into K2EDTA tubes, immediately placed on ice, and centrifuged (5000 rpm, 5 min, 4 °C) to obtain plasma. Brains were snap-frozen in liquid nitrogen. All samples were stored at −80 °C until analysis.

### LC–MS/MS Quantification of EdU in Plasma and Brain

For sample preparation, brains were weighed and diluted 1:2 with phosphate-buffered saline (PBS), then homogenized using zirconium oxide beads in 2 mL tubes at 5000 RPM on a Precellys 24 homogenizer (VWR Bead Mill Homogenizer), applied twice for 20 seconds each.

Plasma (50 µL) or brain homogenate (diluted 1:1 with control plasma) was precipitated with methanol containing D3-methotrexate (100 ng/mL) as the internal standard. After vortexing and centrifugation (10,000 RPM, 10 min, 4 °C), 50 µL of the supernatant was diluted with 100 µL of water for injection.

Liquid chromatography was performed on an Agilent 1260 Infinity II system equipped with a Waters XSelect HSS T3 column (50 × 2.1 mm, 3.5 µm) at 30 °C. The mobile phases were (A) water with 0.1% formic acid and (B) methanol with 0.1% formic acid. The gradient was 10% B (0– 1.0 min) to 50% B (3.0 min), returning to 10% B for a total run time of 7.0 min, at a flow rate of 0.2 mL/min. Injection volume was 4 µL.

Mass spectrometric detection used an Agilent 6470 triple quadrupole with AJS-ESI in positive-ion MRM mode. The following transitions were monitored: m/z 253→136.9 (plasma) and 253→117 (tissue) for EdU, and m/z 458.2→311.2 for the internal standard. Quantification was performed using peak-area ratios relative to matrix-matched calibration standards, following method validation.

### Pharmacokinetic analysis

PK parameters were determined by non-compartmental analysis as described previously (61). Maximum plasma concentration (C_max_) and time to maximum concentration (t_max_) were derived directly from concentration-time profiles. The area under the concentration–time curve from 0 to 24 hours (AUC_0-24h_) was calculated using the linear trapezoidal rule. The terminal elimination rate constant (kel) was obtained from the log-linear portion of the plasma concentration–time curve, and the elimination half-life (t_1/2_) was calculated as ln2/k_el._ The area under the curve extrapolated to infinity (AUC_0–∞_) was determined as AUC_0–24h_ + (C_24h/kel_). Apparent clearance (CL/F) was calculated as Dose/AUC_total_, and apparent volume of distribution (Vd/F) as (CL/F)/k_el_.

## Acknowledgements

We wish to thank UNC Pathology Services Core (PSC) and **Nicholas Pankow** in the UNC PSC for expert technical assistance with Histopathology and Digital Pathology. The UNC PSC is supported in part by an NCI Center Core Support Grant (P30CA016086). We also thank **Dr. Rani Suzanne Sellers** for expert assistance with toxicity scoring of peripheral organs and histopathologic review of brain and peripheral organs. All experiments using passage-zero patient tumor tissues was completed within the Screening Live Cancer Explants Core Facility at UNC. This research is supported by the National Center for Advancing Translational Sciences through NC TraCS (K12TR004416), and National Institute of Health (NIH) Grants ES033414 and GM118102. H.K. is supported by Fulbright PhD Grant. We would like to also extend our thanks to UNC Biomedical Research Imaging Center, UNC Advanced Translational Pharmacology and Analytical Chemistry (ATPAC) Core, UNC Animal Clinical Chemistry Core, and UNC SLiCE Core.

## FIGURE LEGENDS

**Figure S1: Testing the synergism of TMZ+EdU *in vitro* in U87 cell line, extended BLI images and H&E staining on the brains of mice bearing orthotopic xenografts of U87 cells. (A)** Cell survival assay via BLI and CompuSyn assessment of synergy with combination index (CI) values on U87 cell line with the combination of 40 μM EdU (IC50 for U87) and various doses of TMZ (log(CI)<0, log(CI)=0 and log(CI)>0 show synergism, additive effect and antagonism between EdU and TMZ, respectively). Extended BLI images of mice bearing orthotopic xenografts of U87 cells on Days 7 **(B)**, 21 **(C)**, 31 **(D)**, 38 **(E)**, 48 **(F)**, and 58 **(G)**. **(H, I)** H&E staining showing no detectable tumor on the brains of 2 surviving mice from the 5 mg/kg TMZ + 200 mg/kg EdU treatment group.

**Figure S2: Testing the synergism of TMZ+EdU *in vitro* in GBM8 cell line, extended BLI images of mice bearing orthotopic xenografts of GBM8 Cells. (A)** Cell survival assay via BLI and CompuSyn assessment of synergy with combination index (CI) values on GBM8 cell line with the combination of 50 nM EdU (IC50 for GBM8) and various doses of TMZ (log(CI)<0, log(CI)=0 and log(CI)>0 show synergism, additive effect and antagonism between EdU and TMZ, respectively). Extended BLI images of mice bearing orthotopic xenografts of GBM8 cells on Days 9 **(B)**, 42 **(C)**, 94 **(D)**, 133 **(E)**, 150 **(F)**, and 168 **(G)**.

**Figure S3: Spaghetti plots of bioluminescence imaging (BLI) signal fold changes of the tumors of mice bearing orthotopic xenografts of GBM8 and treated with either PBS (n=5) (A)**, 200 mg/kg EdU (n=6) **(B)**, 5 mg/kg TMZ (n=7) **(C)**, 5 mg/kg TMZ + 200 mg/kg EdU (n=7) **(D)**, 1 mg/kg TMZ (n=6) **(E)**, or 1 mg/kg TMZ + 200 mg/kg EdU (n=7) **(F)**.

**Figure S4: Testing the synergism of TMZ+EdU *in vitro* in LN229 cell line and extended BLI images of mice bearing orthotopic xenografts of LN229 Cells. (A)** Cell survival assay via BLI and CompuSyn assessment of synergy with combination index (CI) values on LLN229 cell line with the combination of 20 μM EdU (IC50 for LN229) and various doses of TMZ (log(CI)<0, log(CI)=0 and log(CI)>0 show synergism, additive effect and antagonism between EdU and TMZ, respectively). Extended BLI images of mice bearing orthotopic xenografts of GBM8 cells on Days 4 **(B)**, 24 **(C)**, 64 **(D)**, 70 **(E)**, 73 **(F)**, and 98 **(G)**.

**Figure S5: Toxicity profiling of the mice bearing orthotopic xenografts of U87 cells.** Summary of toxicity findings on the peripheral organs (small intestine, kidneys, spleen, and liver) of mice bearing orthotopic xenografts of U87 cells and treated with either PBS, 200 mg/kg EdU, 5 mg/kg TMZ, or 5 mg/kg TMZ + 200 mg/kg EdU. Abbreviations: 200 mg/kg EdU abbreviated as “EdU”, 5 mg/kg TMZ abbreviated as “TMZ”, 5 mg/kg TMZ + 200 mg/kg EdU abbreviated as “TMZ+EdU”. Grading: 0=no finding; 1=minimal finding; 2=mild finding; 3=moderate finding; 4=marked finding; 5=severe finding.

**Figure S6: Mean body weight and mean body weight change of the mice bearing orthotopic xenografts of U87 (A)**, GBM8 (B), or LN229 (C) cells.

**Figure S7: Complete blood counts for mice bearing orthotopic xenografts of LN229 cells treated with either PBS, 200 mg/kg EdU, 5 mg/kg TMZ, 5 mg/kg TMZ + 200 mg/kg EdU, 1 mg/kg TMZ, or 1 mg/kg TMZ + 200 mg/kg EdU on Days 16, 30 and 58. Complete counts of white blood cells (WBCs) (A)**, lymphocytes (LYMPH) **(B)**, neutrophils **(C)**, basophils (BASO) **(D)**, eosinophils (EO) **(E)**, and red blood cells (RBC) **(F)** for mice bearing orthotopic xenografts of LN229 cells treated with either PBS, 200 mg/kg EdU, 5 mg/kg TMZ, 5 mg/kg TMZ + 200 mg/kg EdU, 1 mg/kg TMZ, or 1 mg/kg TMZ + 200 mg/kg EdU on Days 16, 30 and 58. Lower limit of 95% CI of mean blood counts for female BALB/c nude mice (n=136): White blood cells: 2.97 K/μL, lymphocytes: 1.25 K/μL, neutrophils: 1.02 K/μL, basophils: 0.00 K/μL, eosinophils: 0.01 K/μL, and red blood cells: 8.87 M/μL. One-way ANOVA, *p<0.05. Abbreviations: 200 mg/kg EdU abbreviated as “EdU”, 5 mg/kg TMZ abbreviated as “TMZ (H)”, 5 mg/kg TMZ + 200 mg/kg EdU abbreviated as “TMZ (H) + EdU”, 1 mg/kg TMZ abbreviated as “TMZ (L)”, and 1 mg/kg TMZ + 200 mg/kg EdU abbreviated as “TMZ (L) + EdU”.

## REFERENCES

1. Stupp R, Mason WP, van den Bent MJ, Weller M, Fisher B, Taphoorn MJB, et al. Radiotherapy plus concomitant and adjuvant temozolomide for glioblastoma. N Engl J Med. 2005 Mar 10;352(10):987–96.

2. Rong L, Li N, Zhang Z. Emerging therapies for glioblastoma: current state and future directions. J Exp Clin Cancer Res. 2022 Apr 15;41(1):142.

3. Singh S, Dey D, Barik D, Mohapatra I, Kim S, Sharma M, et al. Glioblastoma at the crossroads: current understanding and future therapeutic horizons. Signal Transduct Target Ther. 2025 Jul 9;10(1):213.

4. Sipos D, Raposa BL, Freihat O, Simon M, Mekis N, Cornacchione P, et al. Glioblastoma: Clinical Presentation, Multidisciplinary Management, and Long-Term Outcomes. Cancers (Basel). 2025 Jan 5;17(1).

5. Haque A, Banik NL, Ray SK. Molecular alterations in glioblastoma: potential targets for immunotherapy. Prog Mol Biol Transl Sci. 2011;98:187–234.

6. Radaelli E, Ceruti R, Patton V, Russo M, Degrassi A, Croci V, et al. Immunohistopathological and neuroimaging characterization of murine orthotopic xenograft models of glioblastoma multiforme recapitulating the most salient features of human disease. Histol Histopathol. 2009 Jul;24(7):879–91.

7. Davis ME. Glioblastoma: overview of disease and treatment. Clin J Oncol Nurs. 2016 Oct 1;20(5 Suppl):S2–8.

8. Hanif F, Muzaffar K, Perveen K, Malhi SM, Simjee SU. Glioblastoma Multiforme: A Review of its Epidemiology and Pathogenesis through Clinical Presentation and Treatment. Asian Pac J Cancer Prev. 2017 Jan 1;18(1):3–9.

9. Goenka A, Tiek D, Song X, Huang T, Hu B, Cheng S-Y. The many facets of therapy resistance and tumor recurrence in glioblastoma. Cells. 2021 Feb 24;10(3).

10. Silver DJ, Lathia JD. Therapeutic injury and tumor regrowth: tumor resection and radiation establish the recurrent glioblastoma microenvironment. EBioMedicine. 2018 May;31:13–4.

11. Rapp M, Baernreuther J, Turowski B, Steiger H-J, Sabel M, Kamp MA. Recurrence pattern analysis of primary glioblastoma. World Neurosurg. 2017 Jul;103:733–40.

12. Jiang H, Yu K, Li M, Cui Y, Ren X, Yang C, et al. Classification of progression patterns in glioblastoma: analysis of predictive factors and clinical implications. Front Oncol. 2020 Nov 3;10:590648.

13. Kawauchi D, Ohno M, Honda-Kitahara M, Miyakita Y, Takahashi M, Yanagisawa S, et al. Clinical characteristics and prognosis of Glioblastoma patients with infratentorial recurrence. BMC Neurol. 2023 Jan 7;23(1):9.

14. Yoon N, Kim H-S, Lee JW, Lee E-J, Maeng L-S, Yoon WS. Targeted genomic sequencing reveals different evolutionary patterns between locally and distally recurrent glioblastomas. Cancer Genomics Proteomics. 2020;17(6):803–12.

15. Wang L, Jung J, Babikir H, Shamardani K, Jain S, Feng X, et al. A single-cell atlas of glioblastoma evolution under therapy reveals cell-intrinsic and cell-extrinsic therapeutic targets. Nat Cancer. 2022 Dec 20;3(12):1534–52.

16. Chehade G, Lawson TM, Lelotte J, Daoud L, Di Perri D, Whenham N, et al. Long-term survival in patients with IDH-wildtype glioblastoma: clinical and molecular characteristics. Acta Neurochir (Wien). 2023 Apr;165(4):1075–85.

17. Lindsey-Boltz LA, Sancar A. New biochemical approaches for treatment of glioblastoma. J Biol Chem. 2025 Sep 20;110748.

18. Fantoni NZ, El-Sagheer AH, Brown T. A Hitchhiker’s Guide to Click-Chemistry with Nucleic Acids. Chem Rev. 2021 Jun 23;121(12):7122–54.

19. Salic A, Mitchison TJ. A chemical method for fast and sensitive detection of DNA synthesis in vivo. Proc Natl Acad Sci USA. 2008 Feb 19;105(7):2415–20.

20. Zeng C, Pan F, Jones LA, Lim MM, Griffin EA, Sheline YI, et al. Evaluation of 5-ethynyl-2’-deoxyuridine staining as a sensitive and reliable method for studying cell proliferation in the adult nervous system. Brain Res. 2010 Mar 10;1319:21–32.

21. Maltsev DI, Mellanson KA, Belousov VV, Enikolopov GN, Podgorny OV. The bioavailability time of commonly used thymidine analogues after intraperitoneal delivery in mice: labeling kinetics in vivo and clearance from blood serum. Histochem Cell Biol. 2022 Feb;157(2):239–50.

22. Wang L, Cao X, Yang Y, Kose C, Kawara H, Lindsey-Boltz LA, et al. Nucleotide excision repair removes thymidine analog 5-ethynyl-2’-deoxyuridine from the mammalian genome. Proc Natl Acad Sci USA. 2022 Aug 30;119(35):e2210176119.

23. Ross HH, Rahman M, Levkoff LH, Millette S, Martin-Carreras T, Dunbar EM, et al. Ethynyldeoxyuridine (EdU) suppresses in vitro population expansion and in vivo tumor progression of human glioblastoma cells. J Neurooncol. 2011 Dec;105(3):485–98.

24. Ligasová A, Strunin D, Friedecký D, Adam T, Koberna K. A fatal combination: a thymidylate synthase inhibitor with DNA damaging activity. PLoS ONE. 2015 Feb 11;10(2):e0117459.

25. Kaanoglu H, Adefolaju A, Fraley C, Shobande M, Dasari R, Mann B, et al. Repurposing the DNA Labeling Agent EdU for Therapy against Heterogeneous Patient Glioblastoma. Mol Cancer Ther. 2025 Aug 1;24(8):1213–25.

26. Han J-H, Yoon JS, Chang D-Y, Cho KG, Lim J, Kim S-S, et al. CXCR4-STAT3 Axis Plays a Role in Tumor Cell Infiltration in an Orthotopic Mouse Glioblastoma Model. Mol Cells. 2020 Jun 30;43(6):539–50.

27. Schulz JA, Rodgers LT, Kryscio RJ, Hartz AMS, Bauer B. Characterization and comparison of human glioblastoma models. BMC Cancer. 2022 Aug 3;22(1):844.

28. Faiz M, Acarin L, Castellano B, Gonzalez B. Proliferation dynamics of germinative zone cells in the intact and excitotoxically lesioned postnatal rat brain. BMC Neurosci. 2005 Apr 12;6:26.

29. Sasportas LS, Kasmieh R, Wakimoto H, Hingtgen S, van de Water JAJM, Mohapatra G, et al. Assessment of therapeutic efficacy and fate of engineered human mesenchymal stem cells for cancer therapy. Proc Natl Acad Sci USA. 2009 Mar 24;106(12):4822–7.

30. Wakimoto H, Kesari S, Farrell CJ, Curry WT, Zaupa C, Aghi M, et al. Human glioblastoma-derived cancer stem cells: establishment of invasive glioma models and treatment with oncolytic herpes simplex virus vectors. Cancer Res. 2009 Apr 15;69(8):3472–81.

31. Sullivan JP, Nahed BV, Madden MW, Oliveira SM, Springer S, Bhere D, et al. Brain tumor cells in circulation are enriched for mesenchymal gene expression. Cancer Discov. 2014 Nov;4(11):1299–309.

32. Sesen J, Dahan P, Scotland SJ, Saland E, Dang V-T, Lemarié A, et al. Metformin inhibits growth of human glioblastoma cells and enhances therapeutic response. PLoS ONE. 2015 Apr 13;10(4):e0123721.

33. DepMap Cell Line Summary [Internet]. [cited 2025 Oct 14]. Available from: https://depmap.org/portal/cell_line/ACH-000075?utm_source=chatgpt.com&tab=overview

34. Ishii N, Maier D, Merlo A, Tada M, Sawamura Y, Diserens A-C, et al. Frequent Co-Alterations of TP53, p16/CDKN2A, p14, PTEN Tumor Suppressor Genes in Human Glioma Cell Lines. Brain Pathol. 1999 Jul;9(3):469–79.

35. Liver, Hepatocyte - Glycogen Accumulation and Depletion - Nonneoplastic Lesion Atlas [Internet]. [cited 2025 Oct 14]. Available from: https://ntp.niehs.nih.gov/atlas/nnl/hepatobiliary-system/liver/Hepatocyte-GlycogenAccumulationandDepletion?utm_source=chatgpt.com

36. Fuller RW, Diller ER. Diurnal variation of liver glycogen and plasma free fatty acids in rats fed ad libitum or single daily meal. Metab Clin Exp. 1970 Mar;19(3):226–9.

37. Rothacker DL, Kanerva RL, Wyder WE, Alden CL, Maurer JK. Effects of variation of necropsy time and fasting on liver weights and liver components in rats. Toxicol Pathol. 1988;16(1):22–6.

38. Wells MY, Weisbrode SE, Maurer JK, Capen CC, Bruce RD. Variable hepatocellular vacuolization associated with glycogen in rabbits. Toxicol Pathol. 1988;16(3):360–5.

39. Mann B, Zhang X, Bell N, Adefolaju A, Thang M, Dasari R, et al. A living ex vivo platform for functional, personalized brain cancer diagnosis. Cell Rep Med. 2023 Jun 20;4(6):101042.

40. Satterlee AB, Dunn DE, Lo DC, Khagi S, Hingtgen S. Tumoricidal stem cell therapy enables killing in novel hybrid models of heterogeneous glioblastoma. Neuro Oncol. 2019 Dec 17;21(12):1552–64.

41. Yadav B, Wennerberg K, Aittokallio T, Tang J. Searching for Drug Synergy in Complex Dose-Response Landscapes Using an Interaction Potency Model. Comput Struct Biotechnol J. 2015 Sep 25;13:504–13.

42. Ianevski A, He L, Aittokallio T, Tang J. SynergyFinder: a web application for analyzing drug combination dose-response matrix data. Bioinformatics. 2017 Aug 1;33(15):2413–5.

43. Weller M, Le Rhun E. How did lomustine become standard of care in recurrent glioblastoma? Cancer Treat Rev. 2020 Jul;87:102029.

44. Bristol-Myers Squibb Company. Paraplatin(carboplatin) [package insert] [Internet]. 2010 [cited 2025 Oct 16]. Available from: https://www.accessdata.fda.gov/drugsatfda_docs/label/2010/020452s005lbl.pdf

45. Zeyen T, Treiber A, Schneider M, Potthoff A-L, Schaub C, Roever L, et al. Second-line temozolomide in first recurrent MGMT-methylated glioblastoma after lomustine/temozolomide: Efficacy and safety. Neurooncol Adv. 2025 Apr 28;7(1):vdaf084.

46. Reyhanoglu G, Tadi P. Etoposide. StatPearls. Treasure Island (FL): StatPearls Publishing; 2025.

47. van der Meulen M, Chahal M, Mason WP. The value of etoposide for recurrent glioma. Can J Neurol Sci. 2024 Jul;51(4):509–12.

48. Dhanavath N, Bisht P, Jamadade MS, Murti K, Wal P, Kumar N. Olaparib: A chemosensitizer for the treatment of glioblastoma. Mini Rev Med Chem. 2025;25(5):374– 85.

49. Hanna C, Kurian KM, Williams K, Watts C, Jackson A, Carruthers R, et al. Pharmacokinetics, safety, and tolerability of olaparib and temozolomide for recurrent glioblastoma: results of the phase I OPARATIC trial. Neuro Oncol. 2020 Dec 18;22(12):1840–50.

50. Lesueur P, Lequesne J, Grellard J-M, Dugué A, Coquan E, Brachet P-E, et al. Phase I/IIa study of concomitant radiotherapy with olaparib and temozolomide in unresectable or partially resectable glioblastoma: OLA-TMZ-RTE-01 trial protocol. BMC Cancer. 2019 Mar 4;19(1):198.

51. Kaina B. Temozolomide, procarbazine and nitrosoureas in the therapy of malignant gliomas: update of mechanisms, drug resistance and therapeutic implications. J Clin Med. 2023 Nov 30;12(23).

52. Slika H, Karimov Z, Alimonti P, Abou-Mrad T, De Fazio E, Alomari S, et al. Preclinical models and technologies in glioblastoma research: evolution, current state, and future avenues. Int J Mol Sci. 2023 Nov 14;24(22).

53. Price M, Ballard C, Benedetti J, Neff C, Cioffi G, Waite KA, et al. CBTRUS Statistical Report: Primary Brain and Other Central Nervous System Tumors Diagnosed in the United States in 2017-2021. Neuro Oncol. 2024 Oct 6;26(Supplement_6):vi1–85.

54. Herrlinger U, Tzaridis T, Mack F, Steinbach JP, Schlegel U, Sabel M, et al. Lomustine-temozolomide combination therapy versus standard temozolomide therapy in patients with newly diagnosed glioblastoma with methylated MGMT promoter (CeTeG/NOA-09): a randomised, open-label, phase 3 trial. Lancet. 2019 Feb 16;393(10172):678–88.

55. Chinot OL, Wick W, Mason W, Henriksson R, Saran F, Nishikawa R, et al. Bevacizumab plus radiotherapy-temozolomide for newly diagnosed glioblastoma. N Engl J Med. 2014 Feb 20;370(8):709–22.

56. Jones AB, Tuy K, Hawkins CC, Quinn CH, Saad J, Gary SE, et al. Temozolomide and the PARP Inhibitor Niraparib Enhance Expression of Natural Killer Group 2D Ligand ULBP1 and Gamma-Delta T Cell Cytotoxicity in Glioblastoma. Cancers (Basel). 2024 Aug 15;16(16).

57. Yuan AL, Ricks CB, Bohm AK, Lun X, Maxwell L, Safdar S, et al. ABT-888 restores sensitivity in temozolomide resistant glioma cells and xenografts. PLoS ONE. 2018 Aug 28;13(8):e0202860.

58. Chou T-C, Talalay P. Quantitative analysis of dose-effect relationships: the combined effects of multiple drugs or enzyme inhibitors. Adv Enzyme Regul. 1984 Jan;22:27–55.

59. Chou: ComboSyn Inc; Paramus,(NJ): 2005 - Google Scholar [Internet]. [cited 2025 Oct 12]. Available from: https://scholar.google.com/scholar_lookup?title=ComboSyn%20Inc%20Paramus%20NJ&author=TC%20Chou&author=N%20Martin&publication_year=2005&

60. Chou T-C. Drug combination studies and their synergy quantification using the Chou-Talalay method. Cancer Res. 2010 Jan 15;70(2):440–6.

61. Akyel YK, Seyhan NO, Gül Ş, Çelik M, Taşkın AC, Selby CP, et al. The impact of circadian rhythm disruption on oxaliplatin tolerability and pharmacokinetics in Cry1-/-Cry2-/-mice under constant darkness. Arch Toxicol. 2025 Apr;99(4):1417–29.

