## Supplementary Figures 1-7 for "Combinatorial Treatment of Glioblastoma with Temozolomide (TMZ) Plus 5-Ethynyl-2’-deoxyuridine (EdU)"

**Figure S1**

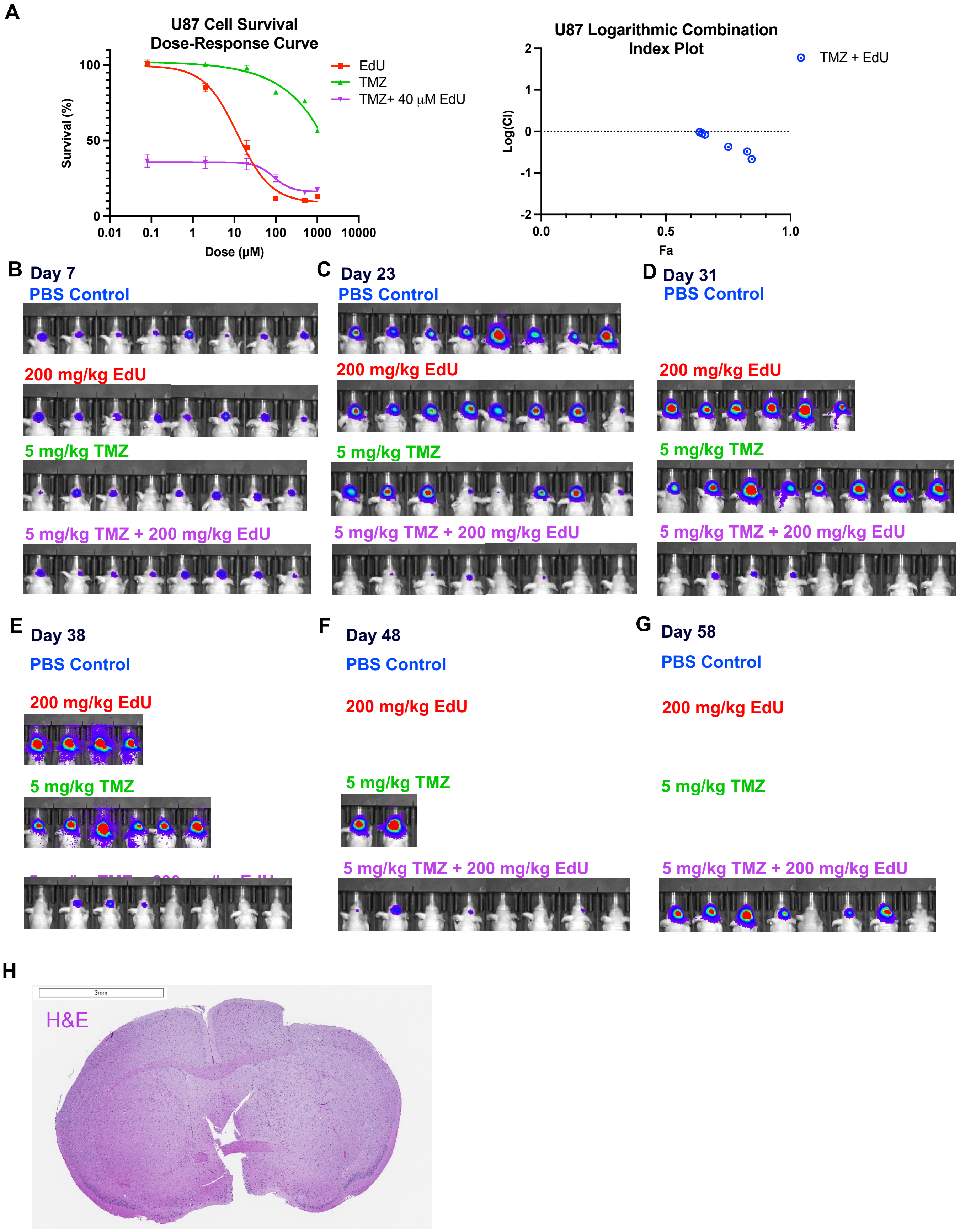
A
U87 Cell Survival
Dose-Response Curve
Survival (%)
Dose (μM)
EdU
TMZ
TMZ+ 40 μM EdU
100
50
0
0.01
0.1
1
10
100
1000
10000
U87 Logarithmic Combination
Index Plot
TMZ + EdU
Log(CI)
Fa
2
1
0
-1
-2
0.0
0.5
1.0
B
Day 7
PBS Control
200 mg/kg EdU
5 mg/kg TMZ
5 mg/kg TMZ + 200 mg/kg EdU
C
Day 23
PBS Control
200 mg/kg EdU
5 mg/kg TMZ
5 mg/kg TMZ + 200 mg/kg EdU
D
Day 31
PBS Control
200 mg/kg EdU
5 mg/kg TMZ
5 mg/kg TMZ + 200 mg/kg EdU
E
Day 38
PBS Control
200 mg/kg EdU
5 mg/kg TMZ
F
Day 48
PBS Control
200 mg/kg EdU
5 mg/kg TMZ
5 mg/kg TMZ + 200 mg/kg EdU
G
Day 58
PBS Control
200 mg/kg EdU
5 mg/kg TMZ
5 mg/kg TMZ + 200 mg/kg EdU
H
3mm
H&E

### Figure S2

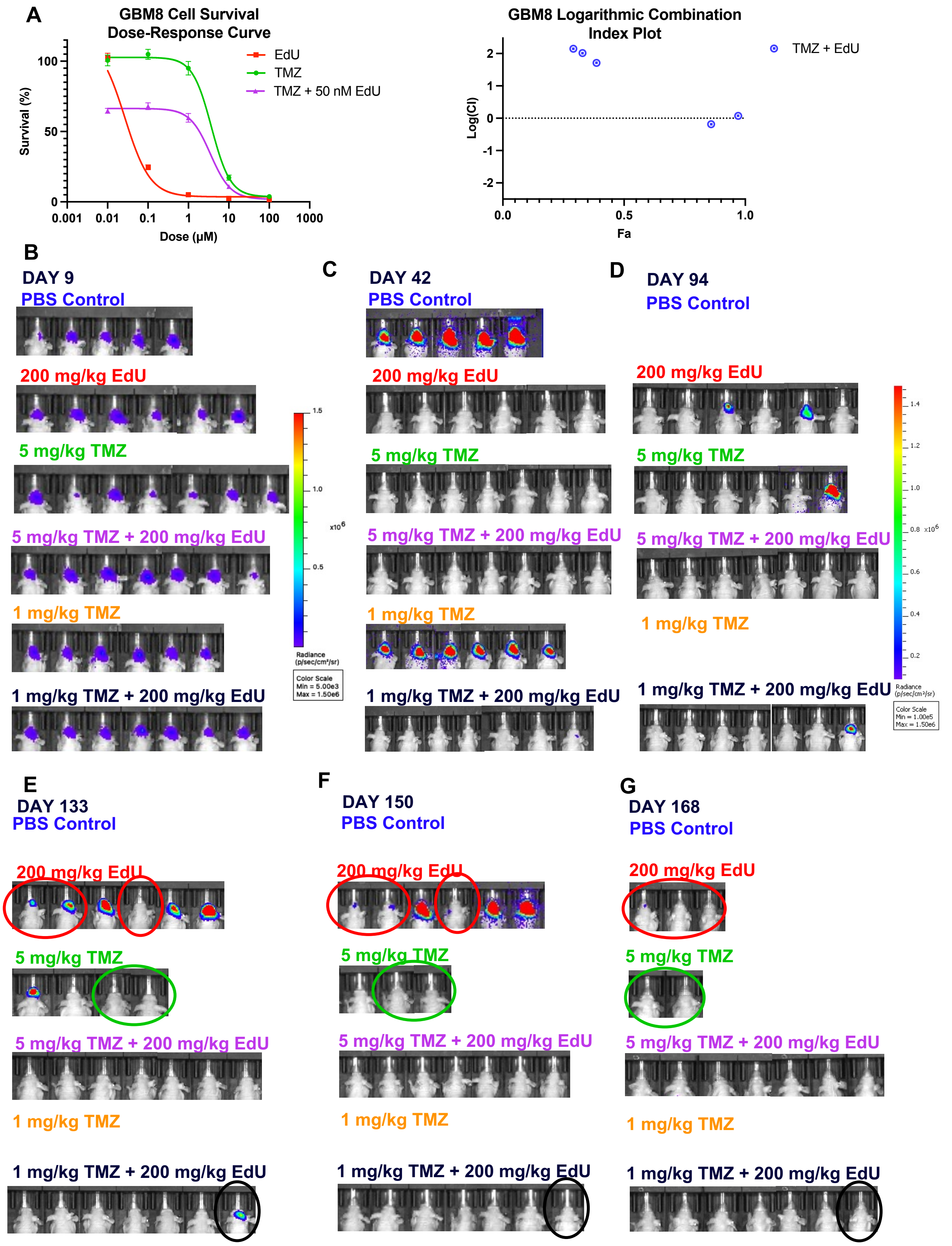
A
GBM8 Cell Survival Dose-Response Curve
Survival (%)
Dose (μM)
0.001
0.01
0.1
1
10
100
1000
0
50
100
EdU
TMZ
TMZ + 50 nM EdU
GBM8 Logarithmic Combination Index Plot
Log(CI)
Fa
0.0
0.5
1.0
TMZ + EdU
B
DAY 9
C
DAY 42
D
DAY 94
PBS Control
200 mg/kg EdU
5 mg/kg TMZ
5 mg/kg TMZ + 200 mg/kg EdU
1 mg/kg TMZ
1 mg/kg TMZ + 200 mg/kg EdU
Radiance (p/sec/cm²/sr)
Color Scale Min = 5.00e3 Max = 1.50e6
Color Scale Min = 1.00e5 Max = 1.50e6
E
DAY 133
F
DAY 150
G
DAY 168

**Figure S3**

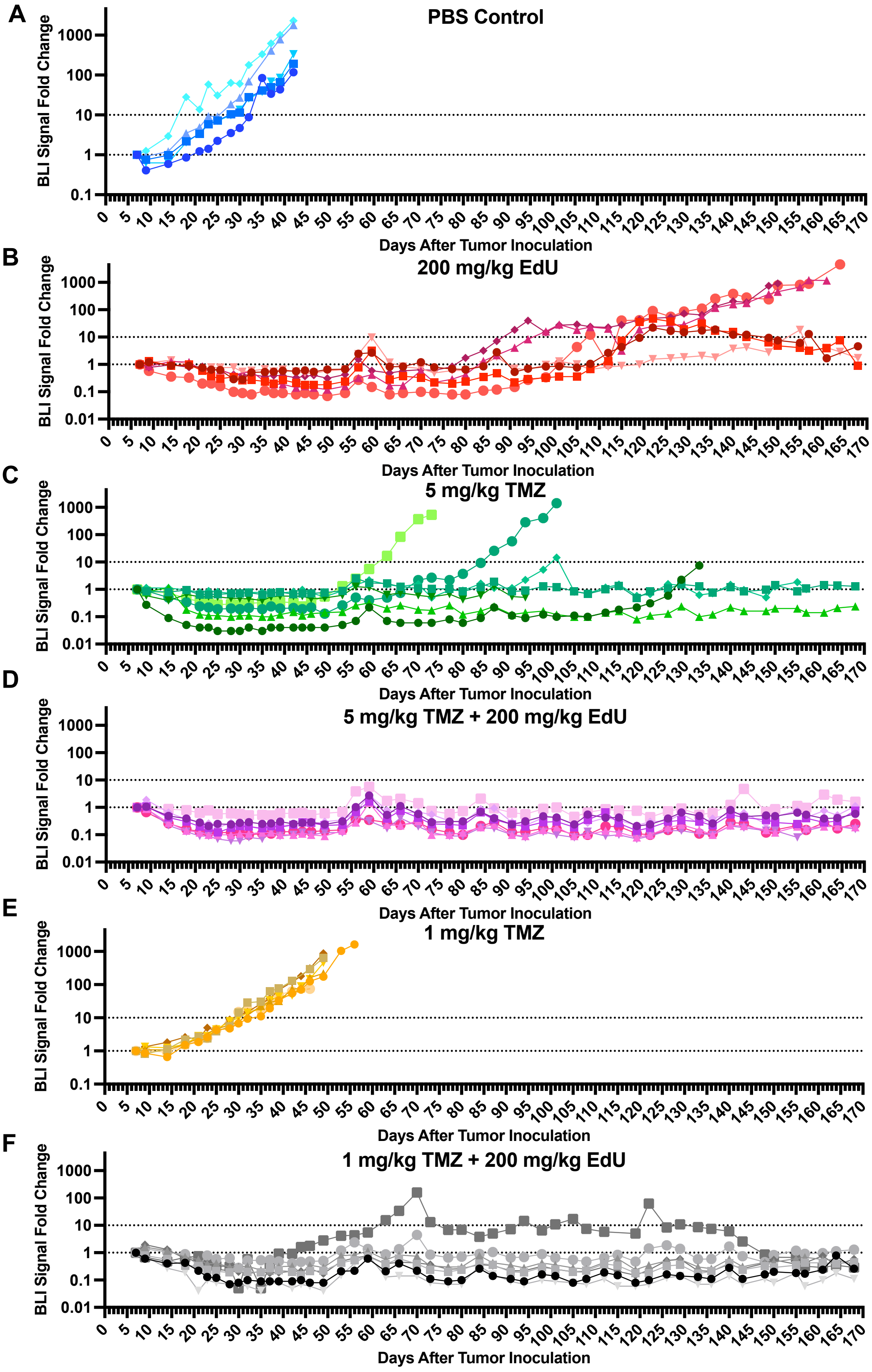

A
PBS Control
B
200 mg/kg EdU
C
5 mg/kg TMZ
D
5 mg/kg TMZ + 200 mg/kg EdU
E
1 mg/kg TMZ
F
1 mg/kg TMZ + 200 mg/kg EdU
BLI Signal Fold Change
Days After Tumor Inoculation

### Figure S4

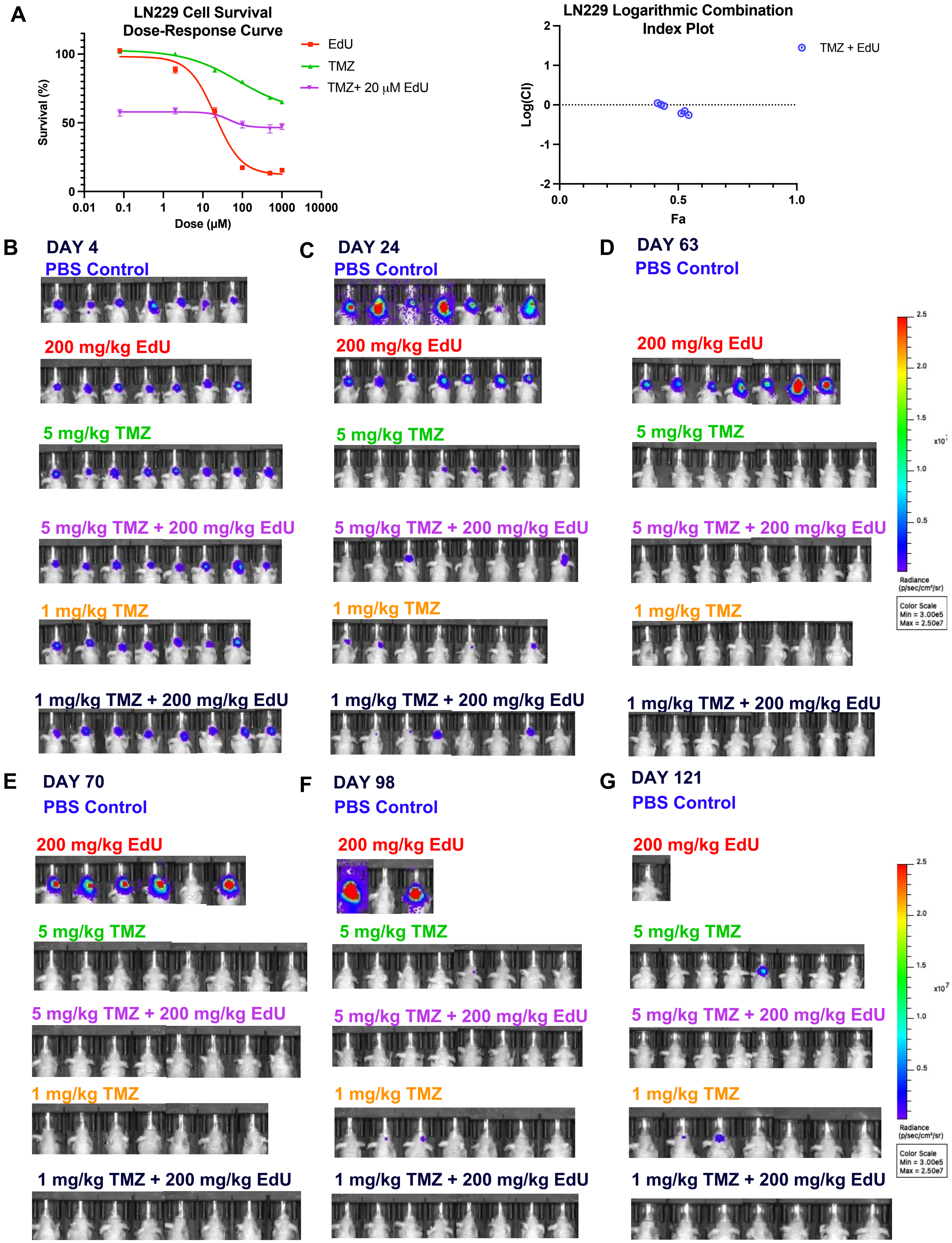

A
LN229 Cell Survival Dose-Response Curve
Survival (%)
Dose (μM)
EdU
TMZ
TMZ+ 20 μM EdU
LN229 Logarithmic Combination Index Plot
Log(CI)
Fa
TMZ + EdU
B
DAY 4
C
DAY 24
D
DAY 63
E
DAY 70
F
DAY 98
G
DAY 121
PBS Control
200 mg/kg EdU
5 mg/kg TMZ
5 mg/kg TMZ + 200 mg/kg EdU
1 mg/kg TMZ
1 mg/kg TMZ + 200 mg/kg EdU
Radiance (p/sec/cm²/sr)
Color Scale
Min = 3.00e5
Max = 2.50e7

### Figure S5

| Study Termination day (D); average for group | 29.1 | 38.5 | 64.8 | 26 | 30.8 |
|---|---|---|---|---|---|
| **Sample ID** | **PBS** | **TMZ** | **TMZ+EdU** | **EdU** | **EdU** |
| ***Microscopic Findings*** | n=7 | n=4 | n=5 | n=6 | n=4 |
| **Small Intestine** | | | | | |
| Single cell necrosis/apoptosis, crypt epithelium | | | | | |
| *minimal* | 0 | 0 | 0 | 3 | 0 |
| *mild* | 0 | 0 | 0 | 1 | 4 |
| Elongated crypts | | | | | |
| *minimal* | 0 | 0 | 0 | 0 | 2 |
| Mitotic figures, increased | | 0 | | | |
| *minimal* | 0 | 0 | 2 | 2 | 2 |
| *mild* | 0 | 0 | 0 | 2 | 0 |

| Study Termination day (D); average for group | 29.1 | 38.5 | 64.8 | 26 | 30.8 |
|---|---|---|---|---|---|
| **Sample ID** | **PBS** | **TMZ** | **TMZ+EdU** | **EdU** | **EdU** |
| ***Microscopic Findings*** | n=7 | n=4 | n=5 | n=6 | n=4 |
| **Kidneys** | | | | | |
| Glomerulopathy, membranoproliferative | | | | | |
| *minimal* | 1 | 0 | 0 | 0 | 0 |
| Tubular degeneration/regeneration | | | | | |
| *minimal* | 0 | 0 | 1 | 1 | 1 |
| Hypertrophy/hyperplasia, cortical tubular epithelium | | | | | |
| *minimal* | 0 | 0 | 0 | 1 | 0 |
| Mineralization, cortical tubular | | | | | |
| *minimal* | 0 | 0 | 1* | 0 | 0 |

| Study Termination day (D); average for group | 29.1 | 38.5 | 64.8 | 26 | 30.8 |
|---|---|---|---|---|---|
| **Sample ID** | **PBS** | **TMZ** | **TMZ+EdU** | **EdU** | **EdU** |
| ***Microscopic Findings*** | n=7 | n=4 | n=5 | n=6 | n=4 |
| **Spleen** | | | | | |
| Increased, extramedullary hematopoiesis* | | | | | |
| *minimal* | 1 | 0 | 1 | 1 | 1 |
| *mild* | 0 | 0 | 3 | 4 | 0 |
| *moderate* | 0 | 0 | 0 | 1 | 0 |

| | | | | | |
|---|---|---|---|---|---|
| **Liver (combined A&B)** | | | | | |
| Vacuolation, hepatocellular, microvacuolar | | | | | |
| *minimal* | 1 | 0 | 1 | 1 | 3 |
| *mild* | 4 | 1 | 3 | 0 | 1 |
| *moderate* | 2 | 1 | 1 | 0 | 0 |
| Vacuolation, hepatocellular, glycogen-like, widespread | | | | | |
| *minimal* | 0 | 1 | 0 | 0 | 0 |
| *mild* | 0 | 0 | 0 | 0 | 1 |
| *moderate* | 0 | 0 | 0 | 6 | 0 |
| *marked* | 0 | 0 | 0 | 0 | 1 |
| Infiltrate, mixed to mononuclear, minimal, multofocal | | | | | |
| *minimal* | 0 | 0 | 1 | 2 | 0 |
| Increased, extramedullary hematopoiesis | | | | | |
| *minimal* | 0 | 0 | 2 | 0 | 0 |
| Kupffer cell cytoplasmic pigment, green | | | | | |
| *minimal* | 0 | 1 | 0 | 0 | 0 |
| Arteriolar hyperplasia | | | | | |
| *Present* | 0 | 1 | 0 | 0 | 0 |

**Figure S6**

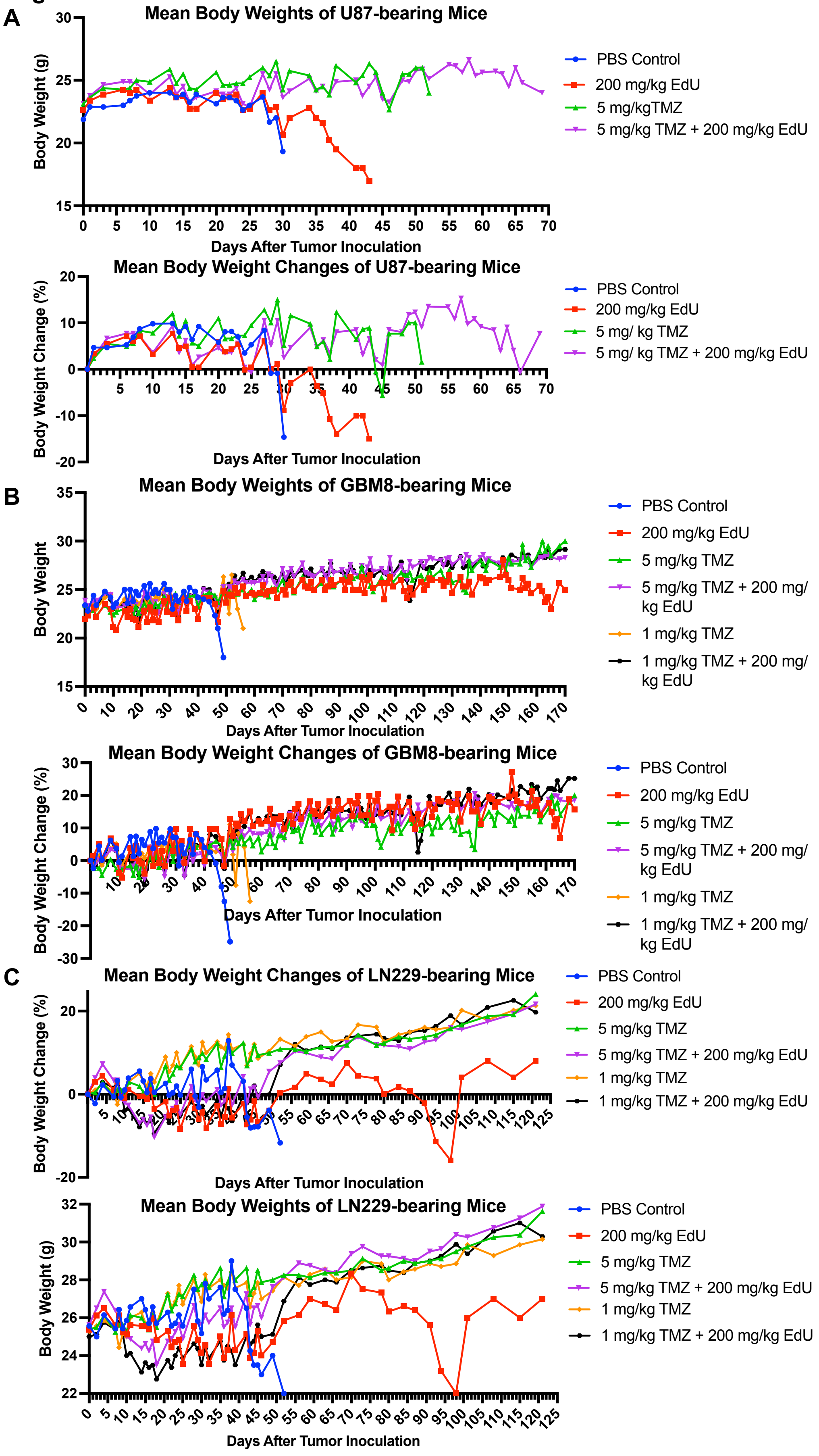
A
Mean Body Weights of U87-bearing Mice
Body Weight (g)
Days After Tumor Inoculation
PBS Control
200 mg/kg EdU
5 mg/kgTMZ
5 mg/kg TMZ + 200 mg/kg EdU
Mean Body Weight Changes of U87-bearing Mice
Body Weight Change (%)
Days After Tumor Inoculation
PBS Control
200 mg/kg EdU
5 mg/ kg TMZ
5 mg/ kg TMZ + 200 mg/kg EdU
B
Mean Body Weights of GBM8-bearing Mice
Body Weight
Days After Tumor Inoculation
PBS Control
200 mg/kg EdU
5 mg/kg TMZ
5 mg/kg TMZ + 200 mg/ kg EdU
1 mg/kg TMZ
1 mg/kg TMZ + 200 mg/ kg EdU
Mean Body Weight Changes of GBM8-bearing Mice
Body Weight Change (%)
Days After Tumor Inoculation
PBS Control
200 mg/kg EdU
5 mg/kg TMZ
5 mg/kg TMZ + 200 mg/ kg EdU
1 mg/kg TMZ
1 mg/kg TMZ + 200 mg/ kg EdU
C
Mean Body Weight Changes of LN229-bearing Mice
Body Weight Change (%)
Days After Tumor Inoculation
PBS Control
200 mg/kg EdU
5 mg/kg TMZ
5 mg/kg TMZ + 200 mg/kg EdU
1 mg/kg TMZ
1 mg/kg TMZ + 200 mg/kg EdU
Mean Body Weights of LN229-bearing Mice
Body Weight (g)
Days After Tumor Inoculation
PBS Control
200 mg/kg EdU
5 mg/kg TMZ
5 mg/kg TMZ + 200 mg/kg EdU
1 mg/kg TMZ
1 mg/kg TMZ + 200 mg/kg EdU

### Figure S7

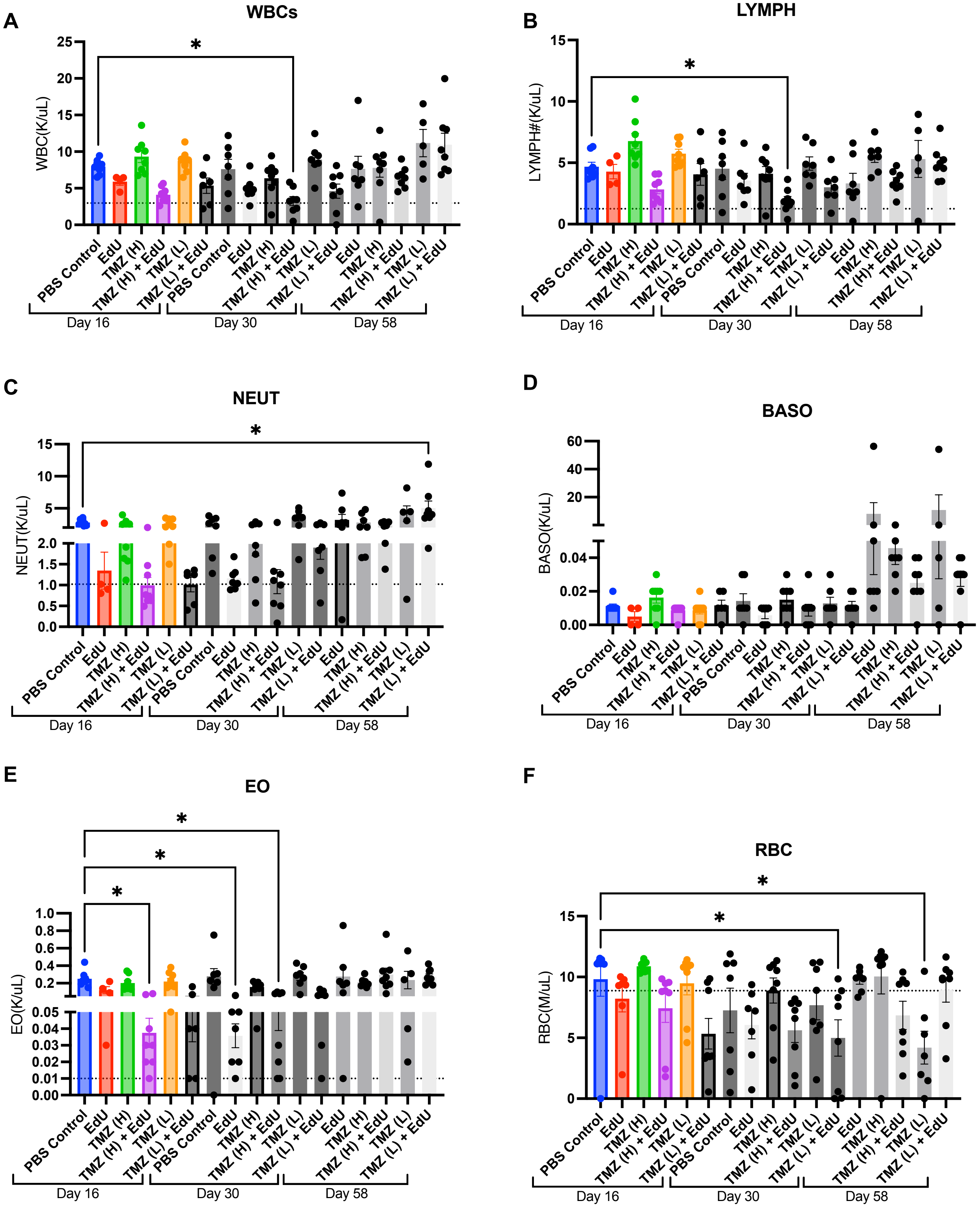
A
WBCs
WBC(K/uL)
0
5
10
15
20
25
*
B
LYMPH
LYMPH#(K/uL)
0
5
10
15
*
C
NEUT
NEUT(K/uL)
0.0
0.5
1.0
1.5
2.0
5
10
15
*
D
BASO
BASO(K/uL)
0.00
0.02
0.04
20
40
60
E
EO
EO(K/uL)
0.00
0.01
0.02
0.03
0.04
0.05
0.2
0.4
0.6
0.8
1.0
*
*
*
F
RBC
RBC(M/uL)
0
5
10
15
*
*
PBS Control
EdU
TMZ (H)
TMZ (H) + EdU
TMZ (L)
TMZ (L) + EdU
TMZ (H)+ EdU
Day 16
Day 30
Day 58
